# Structure-Based Optimization of Pathogen Signal Sequences for Enhanced Antigen Expression in Humans for Vaccine Designs

**DOI:** 10.1101/2025.11.25.690495

**Authors:** Hung N. Do, Apoorv Shanker, Ria Q. Kidner, Makaela M. Montoya, Jessica Z. Kubicek-Sutherland, S. Gnanakaran

## Abstract

Signal peptides (SPs) are short amino acid sequences found at the N-terminus of nascent polypeptides, which serve critical roles in trafficking, folding, and post-translational processing of mature proteins. Many sequence-based computational methods have been developed to predict and design optimal SPs for protein secretion in human cells. In this work, we introduce a structure-based workflow for identifying SPs and their regions, aiming to optimize SPs for protein secretion in human cells. Structural modeling of the SPs in complex with human signal recognition particle 54 kDa (SRP54), combined with hydrophobicity plots of the SPs and identification of the cleavage motif, can be used to detect SPs and SP regions. Afterwards, LigandMPNN is applied to redesign the SPs based on their structural complexes with SRP54. We employ our workflow to optimize the bacterial *Yersinia pestis* F1 SP, Lassa virus glycoprotein (GP) SP, and Venezuelan equine encephalitis virus (VEEV) E3 envelope protein for protein secretion in human cells to serve as vaccine candidates. By comparing the redesigned SPs with the original SPs, we propose that the binding affinity of SPs to SRP54 serve as the most important molecular determinant in the activity of the SPs. Notably, our experiments confirm that the structurally optimized SPs can express the *Y. pestis* F1 protein, Lassa GP, and VEEV GP in human cells. Overall, we demonstrate that structural modeling can serve as a valuable tool to predict the SPs and SP regions for vaccine antigens, predict whether the SPs can express mature proteins in humans, and optimize the SPs for the expression of mature proteins in humans.

## Introduction

Signal peptides (SPs) refer to the short amino acid sequences (typically of 16 to 30 amino acid residues in humans and eukaryotes) that can be found at the N-terminus of nascent polypeptides^1^. These peptides are recognized co-translationally by the signal recognition particle (SRP) and serve to direct proteins to the endoplasmic reticulum (ER) membrane, translocating the nascent chains into the ER lumen to ensure proper trafficking, folding, and post-translational processing of the mature protein^2^.

Human SPs are usually tripartite, which can be divided into three clear regions: n-, h-, and c-regions^3–5^. The n-region, or the N-terminal region, is usually made of one to five residues and populated with positively charged residues to influence membrane topology by interacting with the negatively charged phospholipid head groups in the ER membrane^3,6–8^. The h-region, which could also be referred to as the hydrophobic core, usually includes seven to 15 hydrophobic amino acid residues that can form a helix and are responsible for binding to human SRP 54 kDa (SRP54) to halt the translation and direct the complex to the ER membrane^4,6^. Lastly, the c-region, or the cleavage region, is a polar region containing small, uncharged residues, with the consensus A-X-A motif at the -3/-1 position often recognized by the signal peptidase complex (SPC) for cleavage to release the mature proteins^10^. Overall, SPs direct proteins to the ER-Golgi-plasma membrane complex, targeting secreted proteins, integral membrane proteins, and proteins of the ER and Golgi. The n-region regulates the insertion topology, the h-region interacts with SRP54 and integrates with the membrane, and the c-region is cleaved by the SPC^4,11^.

Several computational tools have been developed to predict SPs and design optimized SPs for optimal protein secretion in mammalian and other expression systems^12–15^. SignalP 6.0 is a widely used and the newest deep learning (DL)-based prediction of SPs, SP regions, and their cleavage sites^12^. The DL model was trained on proteins from archaea, Gram-positive bacteria, Gram-negative bacteria, and eukarya for the prediction of SPs in these species^12^. Another tool to predict SPs is called Phobius^13^, which was published in 2007 and uses a specially designed hidden Markov model (HMM) that detects features to make optimal predictions of transmembrane and signal peptide regions in given amino acid sequences^13^. PredSi, published in 2004, employs a position weight matrix approach trained on datasets comprising proteins from eukarya, Gram-positive bacteria, and Gram-negative bacteria to predict SPs for a large number of protein sequences^16^. Meanwhile, DeepSig, a DL convolutional neural network trained on the dataset, also made of proteins from eukarya, Gram-positive bacteria, and Gram-negative bacteria, was released in 2017 to predict SPs from given amino acid sequences^14^. In terms of SP optimization tools, Barash et al. employed an HMM to describe, predict, identify, and generate a strong artificial SP (secrecon) for the expression of secreted proteins in human cells in 2002^5^. Recently, SecretoGen, a generative transformer trained on millions of naturally occurring SPs from diverse organisms, was published to generate SPs conditional on the mature protein sequences and host organisms for protein secretion^17^.

Since all the currently available methods for predicting and generating SPs are sequence-based, we proposed a structure-based workflow to detect the SPs and their regions for optimization of the SPs for secretion of proteins in human cells in this study. Structural modeling of the SPs in complex with SRP54 by Chai-1^18^, combined with the hydrophobicity plots of the SPs and identification of the cleavage motif, can be used to detect the SP and SP regions. Afterwards, LigandMPNN^16,20^ was applied to redesign the SPs based on their structural complexes with SRP54. Notably, we were able to experimentally confirm the enhanced protein expression of bacterial and viral proteins in human cells using the structurally optimized SPs. By comparing the redesigned SPs with the original SPs, we also proposed the important molecular determinants in the activities of human SPs.

## Results

### Structure-based determination of signal peptide regions

We set out to optimize the recognition of SPs by SRP54, and in turn, optimize the SPs for better expression of their mature proteins, with the goal of enhancing vaccine design. Since the SRP54 primarily interacts with the h-region of the SPs in its M domain, with the n- and c-regions being vital for other functions^6^, our first step involved the determination of the n-, h-, and c-region for each SP so that the h-region can be optimized for better recognition by SRP54, while n- and c-region can be kept fixed for other functions. We first employed SignalP 6.0^12^, a popular tool in predicting SPs and their specific regions, and tested the software in predicting the SP regions for the five SPs of Ebola GP, Lassa GP, Igκ, H1 HAWAII, and bacterial *Y. pestis* F1 protein. Out of the five SPs, signalP 6.0^12^ failed to recognize the two viral SPs—of Ebola and Lassa GPs—and in turn to predict their SP regions (**Supplementary Figure 1A-B**). Meanwhile, the software was able to predict the SP and SP regions for the two commonly used vaccine SPs of Igκ and H1 HAWAII, with MET and MK being the n-regions of Igκ and H1 HAWAII, PAQLLFLLLLWL and AILVVLLYTFT being the h-regions of Igκ and H1 HAWAII, and PDTTG and TANA being the c-regions of Igκ and H1 HAWAII, respectively (**Supplementary Figure 1C-D**), and for bacterial *Y. pestis* SP, with MKK being the n-region, ISSVIAIALFGTIAT being the h-region, and ANA (the canonical A-X-A motif for cleavage by SPC^21–24^) being the c-region (**Supplementary Figure 2A**).

Therefore, an alternative approach using structural modeling was developed to determine the n-, h-, and c-regions of the viral SPs (**Figure 1**). Since the h-region of the SPs, which lies in the middle of the SPs, must be recognized by the SRP54 during the secretion of mature proteins, we directly modelled the structural complexes of the viral SPs of Ebola and Lassa with SRP54 using Chai-1^18^ to determine the n-, h-, and c-region of the two viral SPs (**Figures 1** and **2A-B**). For validation purposes, we also modelled the structural complexes of the vaccine SPs of Igκ and H1 HAWAII with SRP54 (**Figures 1** and **2C-D**) and bacterial *Y. pestis* SP with SRP54 and ffh (the equivalent of SRP54 in bacteria) (**Figure 1** and **Supplementary Figure 2B-C**). The structural models of the bacterial SPs in complex with bacterial signal peptidase 1 and human SPC paralog C (SPC-C) were also built using Chai-1^18^ and included in **Supplementary Figure 3**. Meanwhile, the structural models of the viral SPs in complex with SPC-C were included in **Supplementary Figure 4**. Furthermore, we plotted the hydrophobicity of SPs based on the Kyte-Doolittle^25^ hydrophobicity scale, which could serve as an effective tool for the prediction of SP regions, due to the high hydrophobicity of the h-region (**Supplementary Figure 5**).

**Figure 1.**
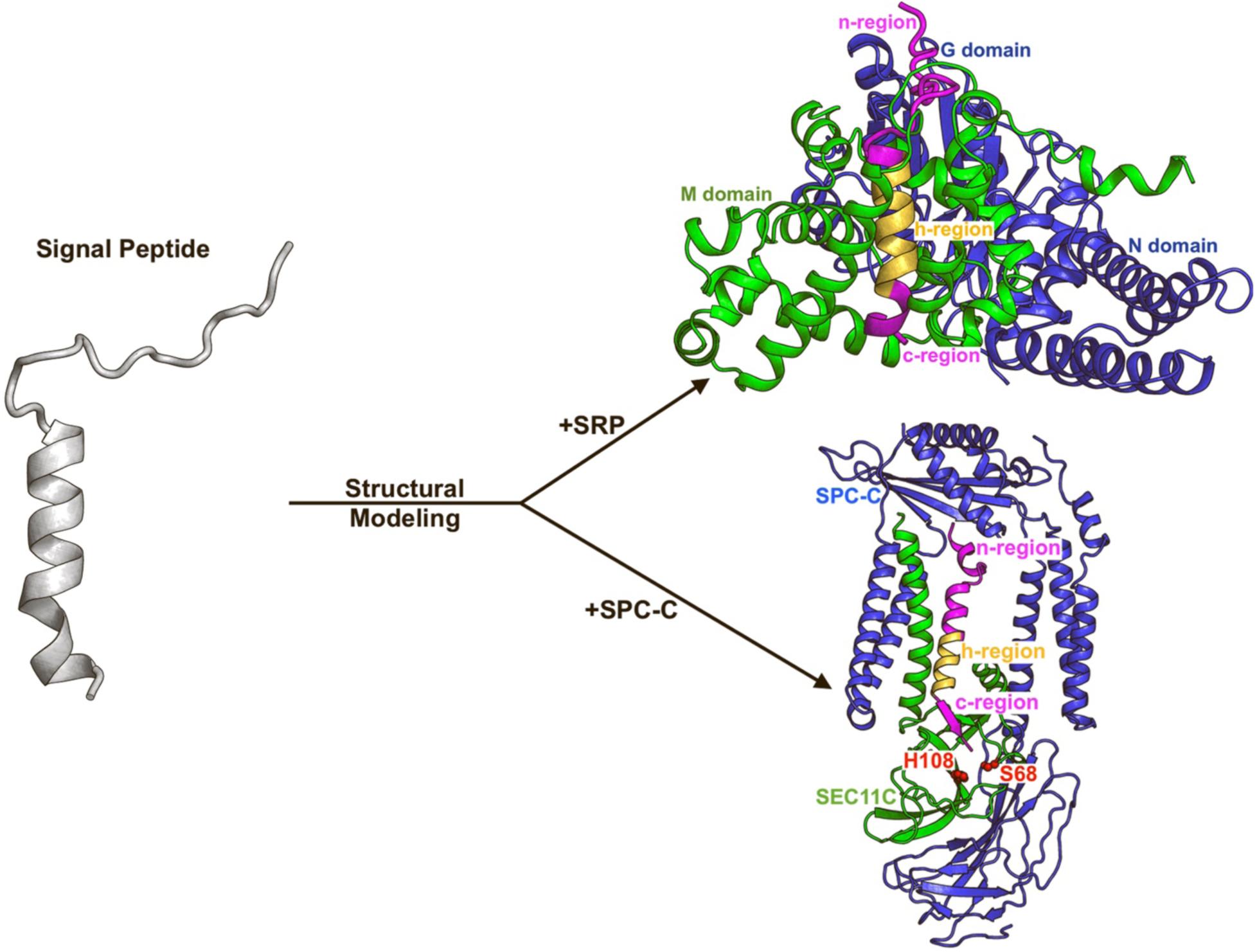
Scheme of the structure-based prediction of signal peptides (SPs) for protein expression in human cells through structural modeling of proteins in complex with the human signal recognition particle 54 kDa (SRP54) and signal peptidase complex paralog C (SPC-C). The approach was particularly useful in predicting the SPs of viral proteins, where signalP6 struggled, to determine the SPs along with their n-, h-, and c-regions.

**Figure 2.**
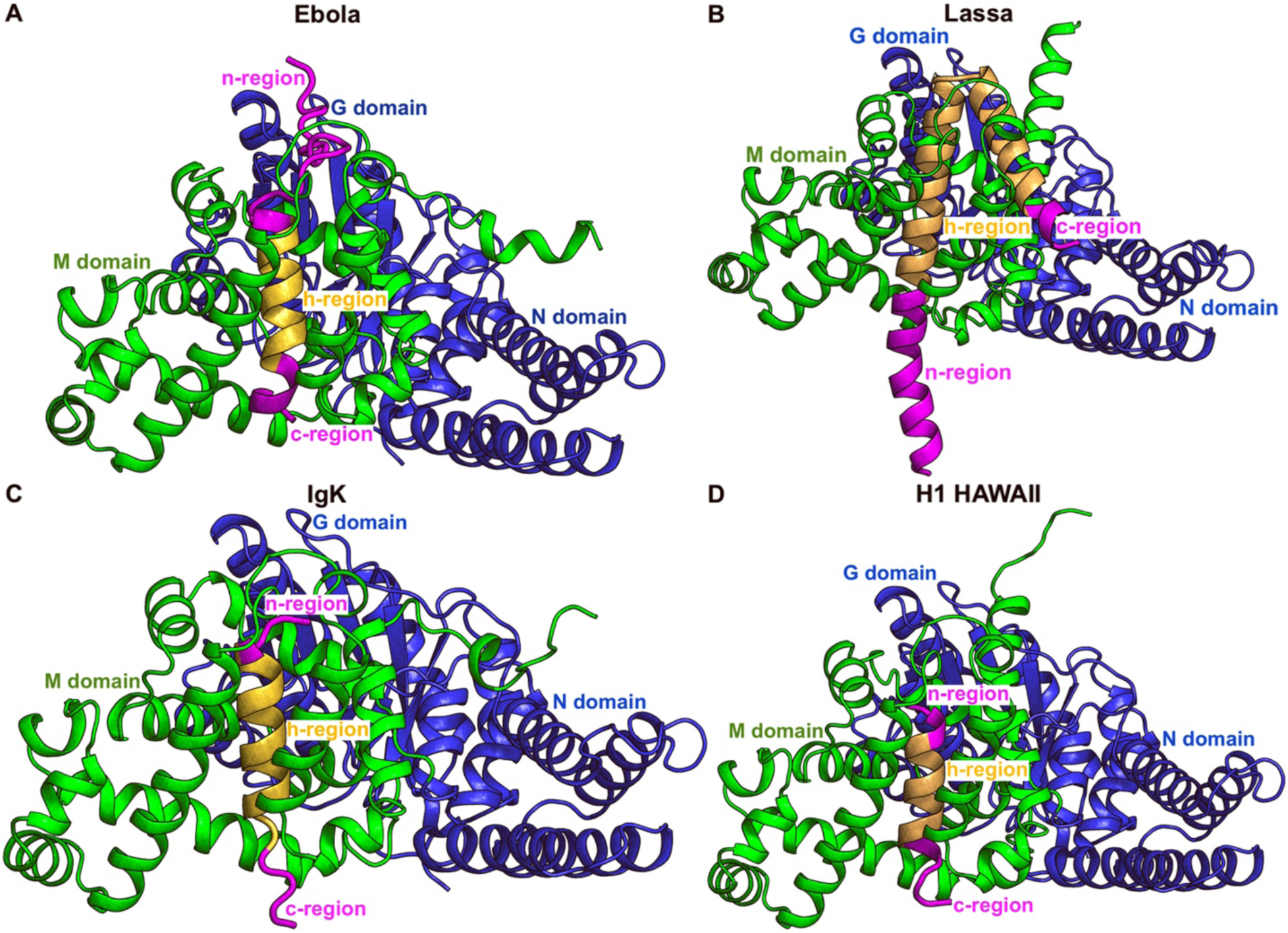
Prediction of the SP regions for the SPs of Ebola and Lassa glycoproteins as well as Igκ and H1 HAWAII (two commonly used SPs for mRNA vaccines) through structural modeling with SRP54. The detailed sequences of the SPs along with their predicted regions through structural modeling were included in **Table 1**.

**Table 1.**
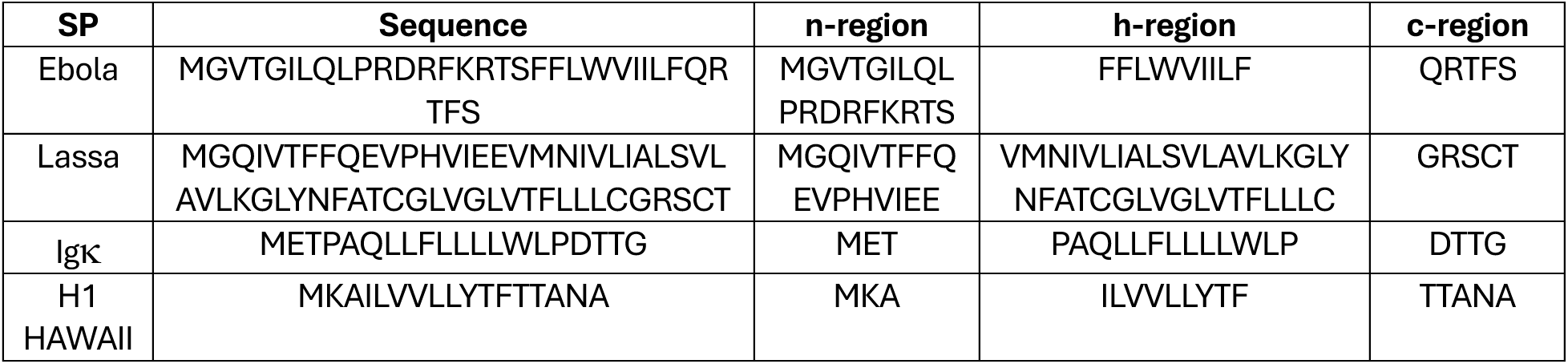
Predicted SP regions for the SPs of glycoproteins and Lassa glycoproteins as well as Igκ and H1 HAWAII SPs through structural modeling with SRP54 as shown in Figure 2.

We deduced the SP regions of the Ebola SP, Lassa SP, Igκ, and H1 HAWAII based on the structural models of their complexes with SRP54 in **Figure 2** and included the results in **Table 1**. Overall, the h-region must exist in helical form and be buried in the pocket formed by the M domain of SRP54^6^. The n-region involved residues located before the h-region, and the c-region involved residues behind the h-region and often ended with the canonical A-X-A or similar motifs including small, uncharged residues, which are favored to be cleaved by the serine protease SPC-C^21–24^. Based on these traits, the h-regions for Ebola SP and Lassa SP were predicted to be FFLWVIILF and VMNIVLIALSVLAVLKGLYNFATCGLVGLVTFLLLC from their respective full sequences of MGVTGILQLPRDRFKRTSFFLWVIILFQRTFS and MGQIVTFFQEVPHVIEEVMNIVLIALSVLAVLKGLYNFATCGLVGLVTFLLLCGRSCT (**Figure 2A-B** and **Table 1**). For Igκ and H1 HAWAII, their predicted h-regions were PAQLLFLLLLWLP and ILVVLLYTF from their respective full sequences of METPAQLLFLLLLWLPDTTG and MKAILVVLLYTFTTANA (**Figure 2C-D** and **Table 1**), being similar to the predictions made by signalP6^12^ (**Supplementary Figure 1C-D**). The prediction for h-region of the bacterial *Y. pestis* SP made from its model complex with ffh and SRP54 (**Supplementary Figure 2B-C**) also agreed with the prediction made by signalP6^12^ (**Supplementary Figure 2A**), with SSVIAIALFGTIAT being the h-region from the full sequence of MKKISSVIAIALFGTIATANA (**Supplementary Figure 2D**). The hydrophobicity plots for Ebola, Lassa, Igκ, H1 HAWAII, and bacterial *Y. pestis* SP showed more positive and consistent hydrophobicity values for the h-regions compared to the n- and c-regions (**Supplementary Figure 5**). In particular, the average hydrophobicity of the h-regions for Ebola, Lassa, Igκ, H1 HAWAII, and bacterial *Y. pestis* SPs was 3.1 ± 1.6, 2.1 ± 2.5, 1.8 ± 2.6, 2.8 ± 2.1, and 2.3 ± 2.1, respectively (**Supplementary Figure 5**). For the n-regions, the average hydrophobicity was -0.4 ± 3.2, 0.3 ± 3.3, -0.8 ± 2.2, -0.1 ± 2.7, and -2.0 ± 2.7, while the average hydrophobicity for the c-regions was -1.3 ± 2.6, -0.8 ± 2.2, -1.3 ± 1.3, -0.3 ± 2.0, and -0.2 ± 2.2, respectively (**Supplementary Figure 5**).

### Molecular determinants in signal peptides for protein expression in human cells

The human serum albumin (HSA) (MKWVTFISLLFLFSSAYS) and H1 HAWAII SPs are two commonly used SPs in mRNA vaccine designs. The HSA SP is native to human, being synthesized by the liver and most abundant protein in human plasma^26^, while H1 HAWAII is non-native to humans and originated from influenza virus (GenBank Protein Accession: QGW43678.1).

We performed structural modeling to characterize the interactions of human serum albumin HSA and H1 HAWAII SP with the endoplasmic reticulum (ER) lipids (consisting of palmitoyloleoylphosphocholine (POPC), palmitoyloleoylphosphoethanolamine (POPE), palmitoyloleoylphosphoinositol (POPI), and palmitoyloleoylphosphoserine (POPS)), human SRP54, and signal peptidase complex paralog C (SPC-C) as well as calculated the hydrophobicity of the two SPs to examine the differences in possible molecular determinants in the two SPs during the classical secretion pathways.

The structural models of H1 HAWAII SP in complex with ER lipids, SRP54, and SPC-C were included in **Supplementary Figure 6A**, **Figure 2D**, and **Supplementary Figure 4D**, while those of HSA SP in complex with ER lipids, SRP54, and SPC-C were shown in **Supplementary Figure 6B**, **6C**, and **6D**, respectively. From the structural modeling with SRP54, we determined the n-region of HSA to be MKW, its h-region to be VTFISLLFLF, and its c-region to be SSAYS (**Supplementary Figure 6C**). The binding energy of the two SPs to the ER lipids, SRP54, and SPC-C were calculated using PRODIGY-LIG^27,28^ for the protein-lipid interactions and PRODIGY^26,30^ for the protein-protein interactions and included in **Supplementary Table 1**. Overall, the contact-based binding energy prediction algorithms of PRODIGY^26,30^ and PRODIGY-LIG^27,28^ predicted lower (i.e., more negative) binding energy of the HSA SP to the ER lipids, SRP54, and SPC-C compared to the H1 HAWAII SP (**Supplementary Table 1**). In particular, the binding energy of the HSA SP to the ER lipids, SRP54, and SPC-C were predicted to be -56.5 kcal/mol, -11.1 kcal/mol, and -10.0 kcal/mol, compared to those of -53.0 kcal/mol, -10.9 kcal/mol, and -6.2 kcal/mol for the H1 HAWAII SP, respectively (**Supplementary Table 1**). Even after we adjusted for the differences in the number of contacts, the binding energy per contacts for HSA binding to SRP54 and SPC-C were still more negative than H1 HAWAII (**Supplementary Table 1**). Specifically, HSA formed 111 contacts with SRP54, and the average binding energy per contact was -0.1 kcal/mol, while H1 HAWAII also formed 111 contacts with SRP54, and the average binding energy contact was -0.098 kcal/mol (**Supplementary Table 1**). Furthermore, HSA formed 71 contacts with SPC-C, averaging -0.14 kcal/mol per contact, compared to H1 HAWAII, which formed 69 contacts with SPC-C and averaged -0.13 kcal/mol per contact (**Supplementary Table 1**).

When it comes to the hydrophobicity of the two SPs, the plots of the hydrophobicity of the H1 HAWAII and HSA SP were included in **Supplementary Figures 5D** and **7**. Overall, the h-region of HSA was more consistently hydrophobic compared to the h-region of H1 HAWAII (**Supplementary Figures 5D** and **7**). Specifically, the h-region of HSA consisted of 10 residues, with a total hydrophobicity of 27.0, an average hydrophobicity of 2.7 per residue, and a standard deviation of 1.8 (**Supplementary Figure 7**). Meanwhile, the h-region of H1 HAWAII was made of nine residues, with a total hydrophobicity of 25.1, an average hydrophobicity of 2.8 per residue, and a standard deviation of 2.1 (**Supplementary Figure 5D**). The h-region of HSA only contained small and uncharged residues (**Supplementary Figure 7**), while the h-region of H1 HAWAII contained one large residue towards the end (i.e., Y), leading to its higher standard deviation (**Supplementary Figure 5D**). Given that the HSA SP is more native to humans than H1 HAWAII^26^, the examined molecular determinants demonstrated that the signal peptides can be optimized for expression of proteins in human cells through the optimization of the h-region for increased consistent hydrophobicity and better interactions with SRP54.

### Structure-based optimization of the bacterial signal peptides for expression in human cells

We set out to optimize the native pathogenic SPs for enhanced expressions of pathogenic proteins in human cells for vaccine designs, given the supporting evidence from above on the determinants of h-region. First, we sought to optimize the bacterial SP for enhanced expression of the mature protein in human cells. Specifically, we aimed to optimize the native *Y. pestis* SP for the enhanced expression of its F1 protein. To optimize the bacterial *Y. pestis* SP, we used the structural model of the SP in complex with SRP54 shown in **Supplementary Figure 2C**. We employed LigandMPNN^16,20^, a deep learning-based protein sequence design tool, to optimize the SPs based on their structural models of the SP complexes with SRP54. We kept the residues at the end of the c-region fixed to avoid affecting their cleavages by SPC-C. The top 10 optimized sequences for *Y. pestis* SP for expression of F1 protein in human cells were included in **Supplementary Table 2**. The top sequence (MKKVLAMLLTLIFLNIATANA) with the highest confidence score of 0.48 from LigandMPNN^16,20^ was used for further comparative analyses with the original sequence, using a similar protocol to compare the HSA and H1 HAWAII SP described above. We built structural models of the original bacterial and optimized *Y. pestis* SP in complex with the ER lipids, SRP54, and SPC-C in **Supplementary Figures 8A**, **2C**, and **3B** for the original bacterial SP and **Supplementary Figure 8B**, **8C**, and **8D** for the optimized SP. Their binding energy was calculated and included in **Supplementary Table 3**. The plots of the hydrophobicity of the original bacterial and optimized *Y. pestis* SPs can be found in **Supplementary Figures 5E** and **9**, respectively.

Overall, the binding energy of the *Y. pestis* SP to the ER lipids and SRP54 improved (i.e., became more negative) after the optimization by LigandMPNN^16,20^ applied on its SRP54 complex (**Supplementary Table 3**). The binding energy with the ER lipids was calculated to reduce from - 54.8 kcal/mol to -56.1 kcal/mol, and the binding energy with SRP54 reduced from -10.0 kcal/mol to -12.4 kcal/mol from the original bacterial to the optimized *Y. pestis* SP (**Supplementary Table 3**). On the other hand, the binding energy with SPC-C increased after the optimization, from -6.5 kcal/mol to -8.7 kcal/mol (**Supplementary Table 3**). However, if we adjusted for the differences in number of contacts, the binding energy per contact between the SP and SPC-C also reduced from the original bacterial to the optimized *Y. pestis* SP (**Supplementary Table 3**). In particular, the bacterial *Y. pestis* SP had 80 contacts with SPC-C, leading to an average of -0.12 kcal/mol per contact, while the optimized *Y. pestis* SP only had 65 contacts with SPC-C, averaging -0.14 kcal/mol per contact (**Supplementary Table 3**). For SRP54, the bacterial *Y. pestis* SP showed 113 contacts, averaging -0.09 kcal/mol per contact, whereas the optimized *Y. pestis* SP illustrated 121 contacts, averaging -0.1 kcal/mol per contact (**Supplementary Table 3**). Furthermore, the hydrophobicity of the h-region of the *Y. pestis* SP increased post-optimization, while the consistency of the hydrophobicity across the h-region remained relatively the same (**Supplementary Figures 5E** and **9**). Seemingly, both h-regions of the bacterial and optimized *Y. pestis* SPs consisted of 14 residues (**Supplementary Figures 5E** and **S9**). However, the hydrophobicity of the h-region increased from 31.5 to 36.3, with the average hydrophobicity per residue increased from 2.3 to 2.6, while the standard deviation only increased slightly from 2.1 to 2.2 due to the introduction of an N residue at position 15 in the h-region of the SP (**Supplementary Figures 5E** and **9**). Overall, it appeared that the optimized *Y. pestis* SP could enhance the expression of its F1 protein due to the lower binding energy to SRP54 and increased hydrophobicity at the h-region following optimization (see experimental results).

### Structure-based optimization of viral signal peptides with post-translational functions

It should be noted that in some cases, such as the Lassa SP and VEEV E3 protein, the SPs remain associated with the mature protein, which has post-targeting functions, including the stabilization of their mature glycoprotein complexes (GPCs) after cleavage^31–34^. Two instances include the Lassa SP for expression of its GPC, and the Venezuelan equine encephalitis virus (VEEV) E3 protein (SLVTTMCLLANVTFPCAQPPICYDRKPAETLAMLSVNVDNPGYDELLEAAVKCPGRKRR), which serves as its SP^31,32^, for expression of VEEV GPC in human cells to serve as a vaccine antigen. While viral SPs can express viral proteins well in human cells (see experimental results), we would like to examine if we could employ a similar structure-based optimization approach to bacterial SPs to redesign the viral SPs with post-translational functions for enhanced protein expression in human cells. Here, we also employed a similar approach with LigandMPNN^16,20^ as we did for the bacterial SP optimization. However, for the Lassa SP and VEEV E3 protein, we also kept the residues involved in the stabilization of their mature GPC fixed, using a contact definition of 4.5 Å between any heavy atoms (i.e., with hydrogen atoms excluded), to ensure the SPs would retain their post-translational functions after optimization.

We built the structural model of three Lassa SPs in complex with its trimeric GPC to determine the SP residues involved in the stabilization, using a combination of Chai-1^18^ and homology modeling using MODELLER^35^ (**Supplementary Figure 10A**). From the complex structure, we determined that Lassa SP residues M1, G2, Q3, I4, T6, F7, Q9, E10, V11, H13, V14, E16, E17, M19, N20, I21, L23, I24, L26, S27, V28, A30, V31, G34, and L35 are involved in the interactions and stabilization of the GPC. Therefore, in redesigning the Lassa SP, we also kept these residues fixed. We employed LigandMPNN^16,20^ on the structural model of Lassa SP in complex with SRP54 included in **Figure 2B**, and the top 10 redesigned sequences generated by LigandMPNN^16,20^ were included in **Supplementary Table 4**. The top sequence (MGQILTFLQEVPHVNEETMNITLIFLSVLAVLNGLAIKYPEGGIGAMNALGLLGRSCT) with the highest confidence score of 0.34 by LigandMPNN^16,20^ was used for the comparative analysis with the original Lassa SP. The structural models of the native Lassa SP in complex with ER lipids, SRP54, and SPC-C were included in **Supplementary Figure 10B**, **Figure 2B**, and **Supplementary Figure 10C**, while the structural models of the redesigned Lassa SP were included in **Supplementary Figure 11B**, **11C**, and **11D**, respectively. We also modelled the structural complexes of three optimized Lassa SPs in complex with its trimeric GPC using Chai-1^18^ to evaluate whether the optimized Lassa SPs still retained its stabilizing post-targeting function (**Supplementary Figure 11A**). The binding energy of the SPs with the GPC, ER lipids, SRP54, and SPC-C was included in **Supplementary Table 5**. The plots of the hydrophobicity of the native and optimized Lassa SPs were included in **Supplementary Figures 5B** and **12**, respectively.

Overall, we saw better binding of the Lassa SP to the GPC and SRP54 and relatively worse binding of the SP to the ER lipids and SPC-C post-optimization (**Supplementary Table 5**). In particular, the binding energy of the Lassa SP to the GPC decreased from -8.2 kcal/mol to -13.5 kcal/mol, and the binding energy to the SRP54 decreased from -14.6 kcal/mol to -18.9 kcal/mol (**Supplementary Table 5**). Accounting for the differences in the numbers of contacts, the average binding energy of the Lassa SP to GPC reduced from -0.09 kcal/mol per contact (with 67 contacts in total for the native Lassa SP) to -0.1 kcal/mol per contact (with 133 contacts in total for the optimized Lassa SP) (**Supplementary Table 5**). Meanwhile, the average binding energy to the SRP54 reduced from -0.08 kcal/mol per contact (with 183 contacts in total for the native Lassa SP) to -0.06 kcal/mol per contact (with 221 contacts in total for the optimized Lassa SP) (**Supplementary Table 5**). However, the binding energy of the Lassa SP to the ER lipids and SPC-C increased from -62.8 kcal/mol to -61.9 kcal/mol and from -14.1 kcal/mol to -13.2 kcal/mol post-optimization (**Supplementary Table 5**). After adjustments for the differences in numbers of contacts, the average binding energy of the native and optimized Lassa SPs to the SPC-C was roughly equal. The native Lassa SP bind to the SPC-C with 146 contacts, averaging -0.1 kcal/mol per contact, while the optimized Lassa SP bind to the SPC-C with 137 contacts, averaging -0.1 kcal/mol per contact (**Supplementary Table 5**). Furthermore, the optimization by LigandMPNN^16,20^ appeared to reduce both the hydrophobicity and consistency of hydrophobicity across the h-region of the Lassa SP (**Supplementary Figures 5B** and **12**). The h-region of the Lassa SP consisted of 36 residues. The total hydrophobicity of the native Lassa SP was calculated to be 74.5, with an average hydrophobicity of 2.1 and standard deviation of 2.5 (**Supplementary Figure 5B**). Meanwhile, the total hydrophobicity of the optimized Lassa SP reduced to 45.6, with an average hydrophobicity reduced to 1.3 and standard deviation increased to 2.8 (**Supplementary Figure 12**). Overall, the Lassa SP carried a very long n-region of 17 residues and long h-region of 36 residues, which were uncharacteristic of human SPs^6^. As the hydrophobicity of the SP h-region decreased post-optimization, the molecular determinant of the h-region hydrophobicity might not be indicative of whether the SP could express proteins in humans. However, the better binding of the redesigned viral SP to SRP54 should be predictive of the expression of the viral proteins in humans.

The SP of VEEV E3 protein also serves a post-translational function to stabilize the mature VEEV GPC^31–34^. Like Lassa, we performed structural modeling of three VEEV E3 proteins in complex with its trimeric GPC to determine the VEEV E3 residues involved in its post-targeting functions (**Supplementary Figure 13A**). From the structural model, we determined that VEEV E3 residues Y23, P27, A28, L31, A32, L34, S35, V38, Y43, D44, L47, E48, A49, V51, and K52 are involved in the interactions with VEEV GPC. We built the structural model of the VEEV E3 protein in complex with SRP54 by Chai-1^18^ (**Supplementary Figure 13C**). We determined that the h-region of VEEV E3 was predicted to be SLVTTMCLLANVTFPCAQPPIC, while the c-region was YDRKPAETLAMLSVNVDNPGYDELLEAAVKCPGRKRR. There appeared to be no n-region in the VEEV E3 protein. We employed LigandMPNN^16,20^ to redesign the VEEV E3 protein starting from the model complex of VEEV E3 and SRP54, and the top 10 sequences generated by LigandMPNN^16,20^ for VEEV E3 were included in **Supplementary Table 6**. The top sequence (TLDTVLSLLSESAFPGAIGISDYIDKPAEHLAYLSKVVDSIGYDEALEASVKDPGRKRR) with the highest confidence score of 0.40 by LigandMPNN^16,20^ was used for the comparative analysis with the native VEEV E3 protein. The structural models of the native VEEV E3 protein in complex with the ER lipids and SPC-C were included in **Supplementary Figure 13B** and **13D**, while the structural models of the redesigned VEEV E3 protein in complex with the VEEV GPC, ER lipids, SRP54, and SPC-C were included in **Supplementary Figure 14A**, **14B, 14C**, and **14D**, respectively. The predicted binding energy of the SPs with the VEEV GPC, ER lipids, SRP54, and SPC-C could be found in **Supplementary Table 7**. The plots of the hydrophobicity of the native and redesigned VEEV E3 proteins were included in **Supplementary Figure 15**.

Overall, the binding energy of VEEV E3 protein with the ER lipids, GPC, and SRP54 increased, while its binding energy with the SPC-C decreased post-optimization (**Supplementary Table 7**). In particular, the binding energy of the native VEEV E3 protein to ER lipids was -51.4 kcal/mol and increased to -48.8 kcal/mol post-optimization (**Supplementary Table 7**). The binding energy of the native VEEV E3 protein with its GPC was calculated to be -6.2 kcal/mol, while the binding of the redesigned VEEV E3 protein with VEEV GPC was found to be -5.4 kcal/mol (**Supplementary Table 7**). With SRP54, the binding energy of the native VEEV E3 protein was -16.5 kcal/mol, compared to a value of -18.2 kcal/mol for the redesigned VEEV E3 protein (**Supplementary Table 7**). Adjusting for the differences in the number of contacts, the average binding energy per contact for native VEEV E3 protein with VEEV GPC was -0.11 kcal/mol (with a total of 57 contacts), and the number for redesigned VEEV E3 protein with VEEV GPC was -0.1 kcal/mol (with a total of 53 contacts) (**Supplementary Table 7**). The average binding energy for native VEEV E3 protein with SRP54 was -0.09 kcal/mol per contact (with a total of 190 contacts), similar to the number for redesigned VEEV E3 protein with SRP54, which was -0.09 kcal/mol per contact (with a total of 209 contacts) (**Supplementary Table 7**). Meanwhile, the binding energy of VEEV E3 protein to the SPC-C decreased from -6.2 kcal/mol to -12.2 kcal/mol post-optimization. However, it should be noted that the structural model of native VEEV E3 protein in complex with SPC-C did not have the protein in the cleavage pocket of the SEC11C catalytic subunit (**Supplementary Figure 13D**), whereas the model of redesigned VEEV E3 protein in complex with SPC-C had the protein in the cleavage pocket of the SEC11C catalytic subunit (**Supplementary Figure 14D**). Here, the hydrophobicity and consistency of hydrophobicity of the h-region of the VEEV E3 protein decreased upon optimization by LigandMPNN^16,20^ (**Supplementary Figure 15**). In particular, the VEEV E3 protein had 23 amino acid residues in its h-region (**Supplementary Figure 15**). The total hydrophobicity of the h-region of the native VEEV E3 protein was 24.1, with an average hydrophobicity of 1.1 per residue and a standard deviation of 2.5 (**Supplementary Figure 15A**). Post-optimization, the total hydrophobicity of the h-region of the redesigned VEEV E3 protein decreased to 16.0, with the average hydrophobicity decreased to 0.7 and standard deviation increased to 2.7 (**Supplementary Figure 15B**). Similar to Lassa, the VEEV E3 protein had an uncharacteristic long h-region of 23 residues and c-region of 37 residues, compared to human SPs^6^. While the SP optimization by LigandMPNN^16,20^ did not improve the hydrophobicity of the h-region, the tool improved the binding affinity of the VEEV E3 protein to the SRP54. Since we also observed enhanced mature protein expressions with redesigned Lassa SP and VEEV E3 protein (see experimental results), we propose that the binding affinity of SPs to SRP54 should serve as the most important molecular determinant in protein expression in humans.

### Cellular protein expression from mRNAs with structurally optimized signal peptide sequences

To experimentally validate the functionality and efficacy of structurally optimized SPs in promoting protein expression, we evaluated protein expression in HEK293T cells following mRNA transfection. Due to their robust growth and high transfection efficiency, HEK263T cells are commonly used for *in vitro* validation of mRNA constructs designed to express vaccine antigens^36^. HEK263T cells were transfected with mRNAs and protein expression was analyzed 24 hours post-transfection. The assay details can be found in the Methods section. To assess the effect of SPs on protein trafficking in the cells, the protein was harvested in three fractions: (i) the supernatant fraction, containing secreted proteins, (ii) the cytosolic fraction, containing intracellular proteins, and (iii) the cell membrane fraction, containing the cell surface-associated proteins. Protein expression was assessed *via* Western blotting as described in the Supplementary Information (**Figure 5A**).

The *Y. pestis* F1 mRNA with a SP sequence optimized for eukaryotic expression showed robust protein expression as shown in **Figure 5B**. Most of the protein was found in the cell membrane fraction, followed by the cytosolic fraction. Very little protein was secreted in extracellular medium. Interestingly, F1 protein was exclusively present as oligomers, a well-known feature of F1 protein^37^. We further tested F1 expression using F1 mRNA with two commonly used SPs, i.e., H1 Hawaii and Igκ, added to the N-terminus. Note, these SP sequences were appended to the N-terminus without removing the native F1 SP sequence. As shown in **Supplementary Figure 16**, the optimized SP resulted in 76% higher protein expression compared to the native F1 SP. Notably, neither the H1 Hawaii nor the Igκ SP yielded in any improvement over the native F1 SP, highlighting the constraints of conventional SPs in driving efficient protein expression. Higher expression of bacterial protein with optimized SP can be attributed to its better interaction with SRP54 due to the lower binding free energy and increased hydrophobicity, thereby validating the design pipeline for structurally optimizing bacterial SP for eukaryotic cellular expression. The structure-based workflow to optimize SPs represents significant advancement in mRNA vaccine design for bacterial pathogens, as it can drive markedly higher antigen expression and consequently amplify the resulting immune response.

Similarly, Lassa GP complex with optimized SP exhibited pronounced protein expression in HEK263T cells. Majority of the expressed protein was found in the cytosolic fraction and included both the GP oligomers as well as GP subunits (GP1 (44 kDa) and GP2 (36 kDa))^38^, as shown in **Figure 5C**. The SP domain on the Lassa GP serves additional role beyond ER targeting, including protein stabilization after cleavage. In optimizing the SP domain, residues critical for GPC stabilization were preserved, which constrained the design space for optimization. Lastly, we investigated protein expression in HEK293T cells transfected with an mRNA construct of the VEEV polyprotein with the redesigned E3 SP domain. As shown in **Figure 5D**, majority of the expressed protein was found in the cell supernatant and in its uncleaved form (E3-E2 (pE2)). Like the Lassa GP, the VEEV E3 protein serves additional functions beyond targeting the nascent protein to the ER membrane. The VEEV E3 protein was also redesigned while keeping the residues involved in glycoprotein stabilization preserved. Successful protein expression for Lassa GP and VEEV polyprotein thus provides the first evidence supporting the validity of the optimization pipeline for complex SP domains with functions extending beyond ER targeting during translation.

## Discussion

In this study, we aimed to investigate structure-based strategies for optimizing the SPs to express viral and bacterial proteins in human cells, thereby serving as potential vaccine antigen candidates. First, we built structural models of the SPs in complex with human SRP54 using Chai-1^18^, from which the n-, h-, and c-regions of the SPs could be determined (**Figure 1**). For SPs with post-translational functions, we also constructed structural models of the SPs in complex with their mature proteins to identify the key residues involved in their post-targeting functions (**Figure 4**). We then employed LigandMPNN^16,20^ to optimize the SPs, primarily at the h-region, with most of the n- and c-regions along with the residues involved in the post-translational functions of SPs kept fixed, based on the structural models of the SPs in complex with SRP54 (**Figures 3-4**). Using the described strategies, we redesigned the bacterial *Y. pestis* SP, Lassa SP, and VEEV E3 proteins and enhanced the expression of the *Y. pestis* F1 protein, Lassa GPC, and VEEV GPC in human cells, aiming to serve as vaccine candidates.

**Figure 3.**
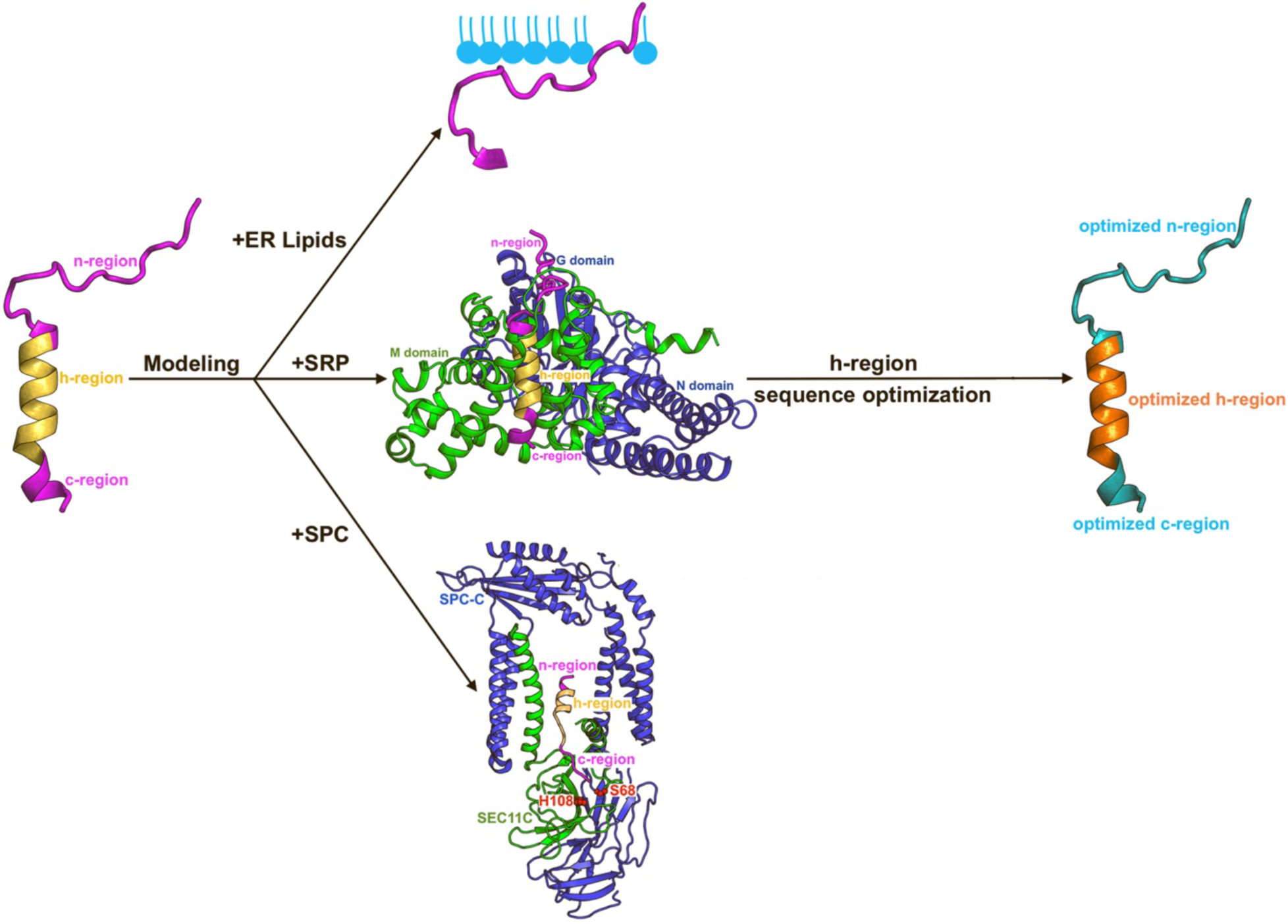
Proposed computational workflow for structure-based optimization of SPs. After the n-, h-, and c-regions of the SPs were determined, structural complexes of the SPs with the n-region interacting with the endoplasmic reticulum (ER) lipids (including phosphatidylcholine PC, phosphatidylethanolamine PE, phosphatidylinositol PI, and phosphatidylserine PS), SPs in complex with the SRP54, and SPs in complex with the signal peptidase complex paralog C (SPC-C) with the c-region in the catalytic subunit SEC11C. The sequences of the h-region of the SPs were then optimized using LigandMPNN^16,20^ to optimize their interactions with the M domain of SRP54, resulting in optimized SPs.

**Figure 4.**
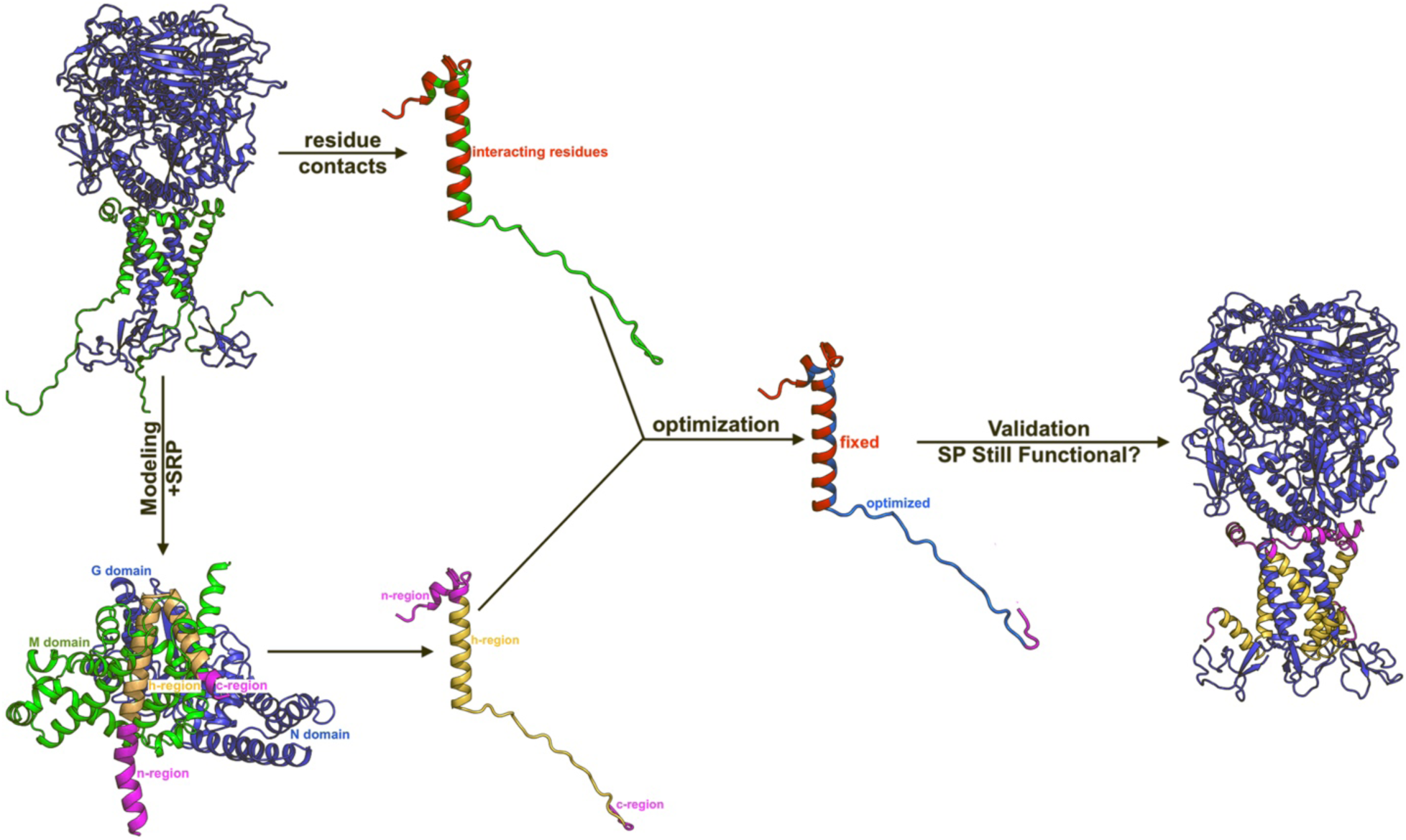
Proposed computational workflow for structure-based optimization of SPs with post-targeting functions. Starting from the complex structures of the SPs bound to the mature proteins, the SP residues involved in the interaction with the mature proteins would be determined. The n-, h-, and c-regions of the SPs could then be determined through the structural modeling with SRP54. The sequences of the functional SPs would then be optimized following the same workflow as in Figure 3, except here the SP residues involved in the interactions with the mature proteins would be kept fixed. After the optimization, the optimized SPs would be validated to determine whether they would still bind to the mature proteins at the proper binding sites and conformations.

**Figure 5.**
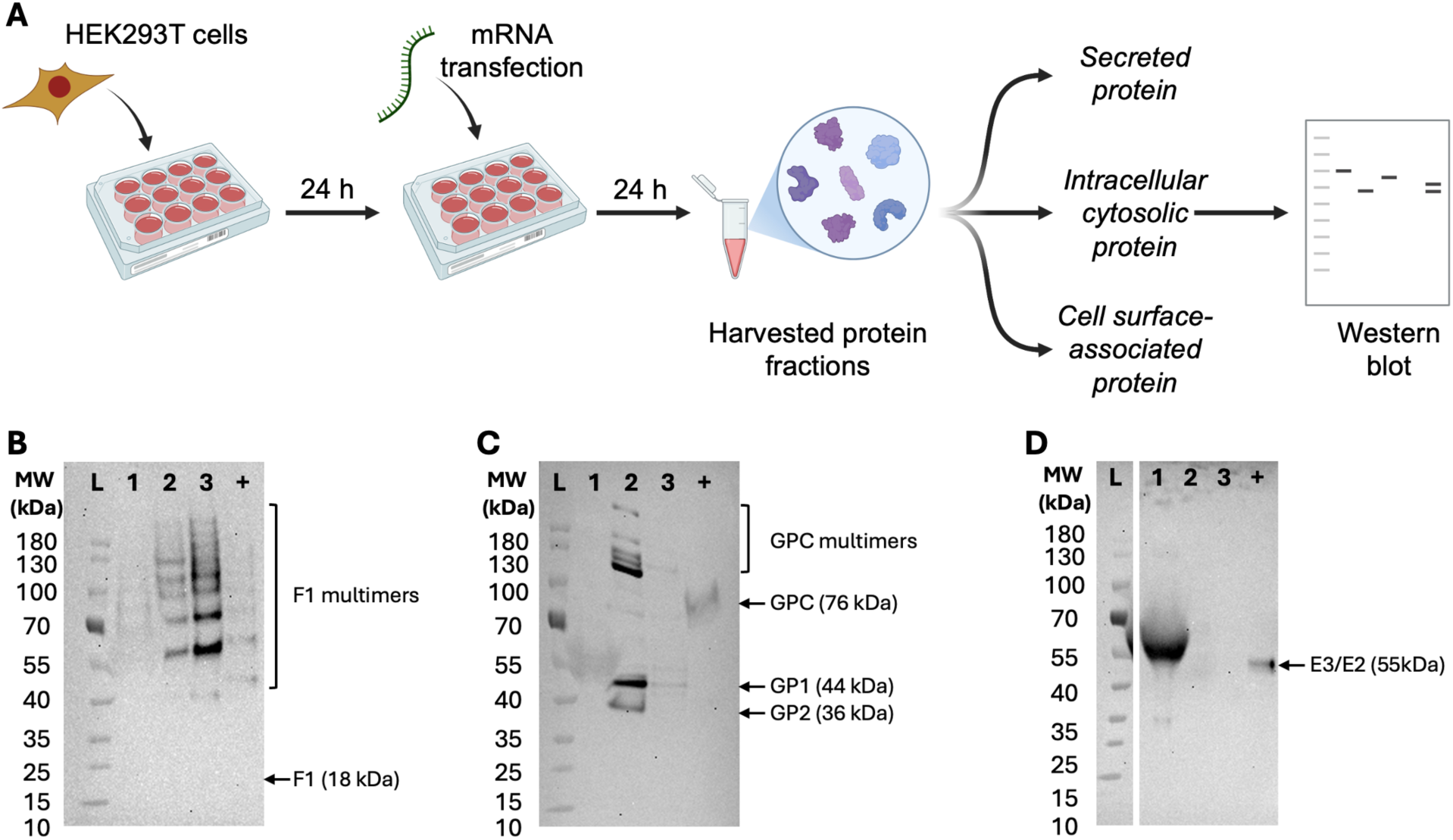
Cellular protein expression following mRNA transfection. **(A)** Schematic overview of the cell transfection assay used to analyze protein expression. **(B)** Western blot analysis of *Y. pestis* F1 protein; Lane designations: L, molecular weight ladder; 1, cell supernatant fraction; 2, cytosolic fraction; 3, cell membrane fraction; +, positive control. **(C)** Western blot analysis of LASV glycoprotein (GPC). **(D)** Western blot analysis of VEEV E3/E2/6k/E1 protein.

We performed structural modeling of the viral and vaccine SPs in complex with human SRP54 to determine the SP regions and found that structural modeling was a good substitute for the cases where SignalP6^12^ failed to predict. From the structural models of the SPs with SRP54, the h-region could be deduced from the helical portions of the SPs located inside the pocket formed by the M domain of SRP54 (**Figure 2**). The n-region consisted of residues prior to the h-region, while the c-region included residues after the h-region and usually ended at the canonical A-X-A (or other small, uncharged residues) motif, which is favored to be cleaved by the serine protease SPC-C^21–24^ (**Figure 2**). The prediction could be verified using the plots of hydrophobicity of the SPs, as there was usually a sudden surge in hydrophobicity at the n/h region boundary and a sudden decrease in hydrophobicity at the h/c region boundary (**Supplementary Figures 5**, **7**, and **G**).

We then used the H1 HAWAII and HSA SPs to evaluate whether a structure-based strategy for optimization of SPs was plausible. HSA is a native SP to human cells^26^, while H1 HAWAII is originated from influenza virus (GenBank Protein Accession: QGW43678.1). We found that the HSA SP bind SRP54 more tightly than the H1 HAWAII SP (**Supplementary Table 1**) and exhibited a more consistent hydrophobicity across its h-region (**Supplementary Figures 5D** and **7**). Here, it should be noted that the contact-based binding energy prediction algorithm of PRODIGY^26,30^ only calculated the binding energy based on the sole input structures. However, proteins are known to be dynamic, so we may carry out molecular dynamics simulations to obtain the most accurate conformations of the SP-SRP54 complexes and employ protocols such as MM-GBSA or MM-PBSA^36^ to calculate the binding energy with biomolecular dynamics accounted for. However, even from the estimated binding energy and hydrophobicity analyses, we could see that HSA bind better to SRP54 than H1 HAWAII due to its higher and more consistent hydrophobicity across the h-region (**Supplementary Figures 5D** and **7**, **Supplementary Table 1)**. Therefore, a structure-based strategy to optimize the SPs for expression of mature proteins in human cells appeared plausible. Furthermore, we hypothesized that the consistency of higher hydrophobicity in the h-region, besides the binding affinity of SPs to SRP54, could serve as an essential molecular determinant in the activity of SPs^40,41^. Later, we also examined whether the propensity of the SP residues in the h-region to form or break helix would play a role in the activity of human SPs (**Supplementary Figure 17**). For HSA versus H1 HAWAII, it appeared that the helical propensity of the h-region was not a determinant, as their proportion of helix-favored residues and helix-breaking residues were similar at 0.3 and 0.1, respectively (**Supplementary Figure 17**).

A deep learning-based protein sequence design tool, LigandMPNN^16,20^, was utilized to optimize the bacterial *Y. pestis* SP, Lassa SP, and VEEV E3 protein for optimal expression of their mature *Y. pestis* F1 protein, Lassa GPC, and VEEV GPC in human cells to serve as vaccine candidates, starting from the structural models of the SPs in complex with human SRP54 (**Supplementary Figure 2C**, **Figure 2B**, and **Supplementary Figure 13C**). Since the bacterial *Y. pestis* SP carried a similar template to human SP, with a short positive n-region, a relatively short hydrophobic n-region, and a small uncharged c-region^10,24,42^, LigandMPNN^16,20^ was able to optimize the *Y. pestis* SP to increase both its binding affinity to the human SRP54 (**Supplementary Table 3**) and the hydrophobicity as well as consistency in hydrophobicity in its h-region (**Supplementary Figures 5E** and **9**). LigandMPNN^16,20^ was also able to increase the helix-favored proportion in the h-region, from 0.3 in the bacterial to 0.7 in the optimized *Y. pestis*, and reduce the helix-breaking proportion in the h-region, from 0.2 to 0.1, of the *Y. pestis* SP post-design (**Supplementary Figure 17**). However, for Lassa SP and VEEV E3 protein, LigandMPNN^16,20^ did not generate sequences that led to increases in hydrophobicity and consistency of hydrophobicity of the h-regions (**Supplementary Figure 5B**, **12**, and **15**) or increases in the helix-favored proportions and decreases in helix-breaking proportions of the h-region (**Supplementary Figure 17**). Nevertheless, the redesigned *Y. pestis* SP, Lassa SP, and VEEV E3 protein were experimentally confirmed to enhance expressions of their mature proteins of *Y. pestis* F1 protein, Lassa GPC, and VEEV GPC (**Figure 5** and **Supplementary Figure 16**). These observations proved to us that the binding affinity of the SPs to SRP54 served as the most important molecular determinant in protein expression in human cells. Therefore, our structure-based optimization algorithm on the native viral SPs could be applied to express other viral proteins in human cells to design vaccines for other pathogens besides those that have been examined in this study.

In conclusion, we have demonstrated that structure-based modeling combined with deep learning-based protein sequence design can serve as a valuable tool to predict the SPs and SP regions for viral and vaccine SPs, determine whether the SPs can facilitate the expression of mature proteins in human cells (i.e., based on the binding poses of the SPs to SRP54), and optimize the SPs for mature protein expression in human cells. We also confirmed that the binding affinity of SPs to SRP54 served as the most important molecular determinant for protein expression in human cells. Our work provides important initial mechanistic insights into designing optimal SPs for the expression of mature proteins in human cells.

## Materials and Methods

### Structure-based determination of signal peptide regions

Since it is necessary for humanized SPs to bind SRP54 to carry out their functions, we set out to optimize the signal peptides by optimizing the hydrophobicity of their h-regions for better interactions with the human SRP54 for optimal expression of vaccine candidates in human cells. We first used signalP6^12^, a popular tool in predicting SPs and their specific regions, to predict the SP regions for the SPs of Ebola and Lassa GPCs, the Igκ and H1 HAWAII SPs (two commonly used SPs to express vaccine proteins in human cells), and the bacterial *Y. pestis* SP for its F1 protein (**Supplementary Figures 1-2** and **Table 1**). SignalP6^12^ failed to predict the SP regions for the first two viral SPs, including Ebola and Lassa (**Supplementary Figure 1A-B**), and was able to predict the SP and SP regions for Igκ, H1 HAWAII, and bacterial *Y. pestis* SP (**Supplementary Figures 1C-D and 2**). Therefore, to determine the SP regions for the viral-like SPs, we performed structural modeling of the SPs in complex with the SRP54 using Chai-1^18^ (**Figure 1**). The h-regions of SPs could be determined from the residues located inside the pocket of the M domain of SRP54, while the n- and c-region would be located outside (**Figure 1**). For confirmation, we also modelled the structures of the SPs in complex with human SPC-C (PDB: 7P2Q)^24^, an important serine protease involved in cleaving the SPs from the mature proteins (**Supplementary Figures 3-4**). Furthermore, we plotted the hydrophobicity of the SPs, calculated based on the Kyte-Doolittle^25^ hydrophobicity scale, of the SPs to further validate the predicted regions of the SPs (**Supplementary Figure 5**).

### Structure-based optimization of signal peptides for protein expression in human cells

It is known that the SPs that are well recognized and bind to the SRP54 could express the proteins of interest well in human cells^40,41^. Therefore, we proposed to employ LigandMPNN^16,20^, a deep learning-based protein sequence design tool, to optimize the sequences of the SPs for better binding and recognition by the SRP54 based on the structural models of SPs in complex with SRP54 generated by Chai-1^18^ (**Figures 3-4**). Throughout this study, LigandMPNN^16,20^ was applied using the “soluble_mpnn” model checkpoint, with the temperature set to 0.05 and random seed set to 111. As the h-region was the primary region in the SP to interact with the SRP54^4^, we chose to only redesign the residues in the SP h-regions and kept the n- and cleavage site of c-regions (usually the last four residues of SPs) fixed (**Figure 3**). For SPs with post-targeting functions, we also kept the residues involved in their post-targeting functions (i.e., binding to the mature proteins) fixed during the sequence optimization using LigandMPNN^16,20^ on their complexes with SRP54 (**Figure 4**). After optimization, we evaluated the recognition of the optimized SPs by SRP54 by constructing the structural models of the optimized SPs in complex with human SRP54 and comparing the binding energy of optimized SPs with the binding energy of the original SPs to the SRP54 to determine if there were any improvements. The binding energy of the two proteins were calculated using PRODIGY^26,30^. We also modelled the structural complexes of the original and optimized SPs with the endoplasmic reticulum (ER) lipids and SPC-C and compared their corresponding binding energy, predicted by PRODIGY-LIG^27,28^ and PRODIGY^26,30^, respectively. We modelled the ER lipids using five molecules of POPC, two molecules of POPE, one molecule of POPI, and one molecule of POPS to mimic the lipid composition of the ER membrane^6–8^. Lastly, we plotted the hydrophobicity of the optimized SPs and compared them to the original SPs to determine if there were improvements in the values and consistency of hydrophobicity (i.e., standard deviation of hydrophobicity for residues) across the h-regions. We validated our proposed computational workflow for structure-based optimization of SPs using the H1 HAWAII and the human serum albumin (HSA) SPs (**Supplementary Figures 6-7** and **Supplementary Table 1**). Afterwards, we employed our proposed computational workflows for structure-based optimization of SPs to optimize the SPs of bacterial *Y. pestis* F1 protein (**Supplementary Figures 8-9** and **Supplementary Tables 2-3**) and Lassa viral GP complex (GPC) (**Supplementary Figures 10-12** and **Supplementary Tables 4-5**) as well as the Venezuelan equine encephalitis virus (VEEV) E3 protein (SLVTTMCLLANVTFPCAQPPICYDRKPAETLAMLSVNVDNPGYDELLEAAVKCPGRKRR) (**Supplementary Figures 13-15** and **Supplementary Tables 6-7**), which serves as the SP for VEEV GPC, to express the respective mature proteins in human cells to serve as vaccine candidates for the respective diseases.

### Cells, materials, and reagents

HEK263T cells (CRL-3216) were procured from American Type Culture Collection (ATCC). Dulbecco’s Modified Eagle Medium (DMEM; D6429), fetal bovine serum (FBS; F242), penicillin-streptomycin (P4333), trypsin-EDTA solution (T4049), Dulbecco’s Phosphate Buffered Saline (DPBS; D8537), bovine serum, albumin (BSA; A9418) and Tween-20 (P1379) were purchased from Millipore Sigma. Cell culture flasks (Corning 3290) and 12-well assay plates (Corning 3512) were purchased from Corning. mRNAs for protein antigens and mLantern green fluorescent protein were supplied by Moderna, Inc. (Cambridge, MA). TransIT-mRNA Transfection Kit (MIR 2250) was purchased from MirusBio. MEM-Per Plus Membrane Protein Extraction Kit (Catalog #86842) and Gibco Opti-MEM reduced serum media (Catalog #31985062) were purchased from ThermoFisher Scientific. The following reagents and supplies for Western blotting were purchased from ThermoFisher Scientific: Bolt 4-12% Bis-Tris Plus WedgeWell (NW04122BOX) and Novex 4-12% Tris Glycine Plus WedgeWell (XP04120BOX) gels, iBlot3 transfer stacks (IB34002), Bolt MOPS SDS Running Buffer (B0001-02), Novex Tricine SDS Running Buffer (LC1675), 10× Bolt Sample Reducing Agent (B0009), Bolt LDS Sample Buffer (4×; B0008), Pierce^TM^ ECL Western Blotting Substrate (32106), and PageRuler Prestained Protein Ladder (10 – 180 kDa; Catalog #26616). Non-fat dry milk powder was purchased from Kroger. SDS-PAGE was performed using an Invitrogen PowerEase Touch HV power supply with an XCell SureLock Mini-Cell electrophoresis system. An Invitrogen iBlot3 system was used to transfer electrophoretically separated proteins from the gel to a nitrocellulose membrane. The membrane was subsequently imaged and analyzed on an Invitrogen iBright 750 gel imaging system. The positive control antigen protein and primary and secondary antibodies used for Western blot analyses were included in **Table 2**.

**Table 2.** Positive control antigen protein and primary and secondary antibodies used for the Western blot analyses shown in Figure 5.

| Antigen | Positive control | Primary antibody | Secondary antibody |
| --- | --- | --- | --- |
| ***Y. pestis* F1** | Recombinant *Y. pestis* F1 antigen protein (His tag) (Abcam ab236940) | Mouse anti-F1 recombinant antibody (clone YPF19; Creative Biolabs MRO-372-MZ) | Goat anti-mouse IgG (H+L), HRP (ThermoFisher #31430) |
| **Lassa LASV GPC** | Lassa glycoprotein from Steven Bradfute (University of New Mexico) | Rabbit anti-LASV GP polyclonal SHMT1 antibody (Abcam ab190655) | Goat anti-rabbit IgG (H+L), HRP (ThermoFisher #32460) |
| **Venezuelan Equine Encephalitis Virus (VEEV) E3/E2/6k/E1** | VEEV E3E2 GP recombinant protein (His-tagged) (IBT Bioservices #0563-001) | Monoclonal Anti-VEEV, PTF-39 (Subtype IB) E2 Glycoprotein Antibody, Clone 1A3B-7 (BEI Resources NR-51619) | Goat anti-mouse IgG (H+L), HRP (ThermoFisher #31430) |

### Cell culture and transfection assay

HEK263T cells were maintained in a DMEM cell growth medium supplemented with 10% FBS and 1% penicillin-streptomycin. For transfection assays, HEK293T cells were seeded in a 12-well assay plate at 1×10^5^ or 2.5×10^5^ cells per well in 1 mL growth medium and incubated for 48 (1×10^5^ cells) or 24 hours (2.5×10^5^ cells) at 37°C and 5% CO_2_. On the day of transfection, cell growth medium was replaced with fresh growth medium lacking penicillin-streptomycin. mRNAs were prepared for transfection using the MirusBio Trans-IT transfection kit according to the manufacturer’s protocol and added to the cells at 2 μg per well. To account for reagent-induced cellular toxicity, one well on the assay plate was treated with the transfection agent alone, without mRNA. As a positive control, cells in two wells were transfected with mLantern mRNA (2 μg per well) to enable visual confirmation of successful transfection. Following mRNA addition, the assay plate was incubated at 37°C and 5% CO_2_ for 24 hours.

After 24 hours, wells transfected with mLantern mRNA were imaged using a fluorescence microscope in the GFP channel (488/525 nm) to confirm transfection efficiency. Protein extraction was performed using a MEM-Per Plus Membrane Protein Extraction Kit, yielding three fractions: (i) the supernatant fraction, containing secreted proteins, (ii) the cytosolic fraction, containing intracellular proteins, and (iii) the cell membrane fraction, containing the cell surface-associated proteins. Extracted proteins were stored at -80°C until further analysis.

### Western blot analysis of protein expression

Expression of protein antigens were confirmed by Western blot analysis using the antibodies listed above. Briefly, the harvested protein fractions were denatured and reduced at 90°C for 10 minutes in an SDS-PAGE buffer containing up to 26 μL of harvested sample or 20 – 100 ng of positive control protein antigens (in 26 μL volume), 10 μL Bolt LDS Sample Buffer, and 4 μL of Bolt Sample Reducing Agent (total volume = 40μL). The samples and the positive control, along with the PageRuler Prestained Protein Ladder, were then loaded onto a Bolt 4-12% Bis-Tris Plus WedgeWell or Novex 4-12% Tris Glycine Plus WedgeWell gel for resolution. Post electrophoretic separation, proteins were transferred to a nitrocellulose membrane using an Invitrogen iBlot3 instrument, according to the manufacturer’s instructions. Blocking and probing of the membrane was performed in 1 DPBS (pH 7.4) with 5% non-fat dry milk or 2.5% BSA and 0.05% Tween-20. Membrane was first blocked by submerging in 15 mL blocking buffer for 30 minutes with shaking. The blocking buffer was then drained and replaced with 15 mL of blocking buffer containing an appropriate dilution of the primary antibody (typically, 1:1000). The membrane was incubated with shaking for an hour at room temperature or overnight at 4°C. The membrane was then washed at least thrice with 1× DPBS containing 0.05% Tween-20 (PBS-T). The membrane was then incubated with shaking for an hour in the secondary antibody solution prepared in the blocking buffer at an appropriate dilution (typically, 1:5000). Subsequently, the membrane was washed extensively with PBS-T and developed in Pierce ECL Western Blotting Substrate for imaging. The membranes were imaged on an Invitrogen iBright 750 gel imaging system.

### Quantification of protein expression

Protein expression levels were quantified from Western blot images using ImageJ (NIH, Bethesda, MD, USA). Gel images were first converted from RGB to 8-bit grayscale format. For each lane, a rectangular region of interest (ROI) encompassing the entire protein band was selected, and the mean gray value was measured. The measured gray values were then inverted by subtracting them from 255 (i.e., 2⁸ − 1). Background correction was performed by subtracting the corresponding values obtained from mock-transfected control lanes for each fraction (cell supernatant, cytosol, and cell membrane).

## Data Availability Statement

Data supporting the findings of this study are included in the article and its Supplementary Information files.

## Code Availability Statement

This study utilized the standard builds of SignalP 6.0^12^ (https://services.healthtech.dtu.dk/services/SignalP-6.0/), Chai-1^18^ (https://github.com/chaidiscovery/chai-lab), PRODIGY^30^ (https://github.com/haddocking/prodigy), PRODIGY-LIG^27^ (https://github.com/haddocking/prodigy-lig), ProteinMPNN^20^ (https://github.com/dauparas/ProteinMPNN), and LigandMPNN^16^ (https://github.com/dauparas/LigandMPNN). All the related parameters were included in the Materials and Methods section.

## Acknowledgements

This work was supported by the Defense Threat Reduction Agency (award no. HDTRA1242031). This work was performed at the Los Alamos National Laboratory (LANL), operated by Triad National Security, LLC, for the National Nuclear Security Administration of the U.S. Department of Energy (contract 86233218CNA000001). Computational resources were provided by LANL Institutional Computing. The authors would like to thank Daphne Stanley for supporting this work. We would also like to thank Moderna for providing the mRNA materials to carry out the in-vitro experiments.

## Author Contributions

H.N.D. performed computational research, analyzed data, and wrote the manuscript. R.Q.K. performed protein extraction and Western blot; M.M. and R.Q.K. performed cell transfection. A.S. supervised research, analyzed data, and wrote the manuscript. J.Z.K. acquired funding, supervised research, and wrote the manuscript. S.G. supervised research, analyzed data, and wrote the manuscript. All authors contributed to the final version of the manuscript.

