## Supplementary Information for "Structure-Based Optimization of Pathogen Signal Sequences for Enhanced Antigen Expression in Humans for Vaccine Designs"

**Supplementary Figure 1. Results of signalP6 predictions of the SPs of the Ebola and Lassa glycoproteins as well as Igκ and H1 HAWAII (two commonly used SPs for mRNA vaccines).** The program failed to predict the SPs for the viral proteins.

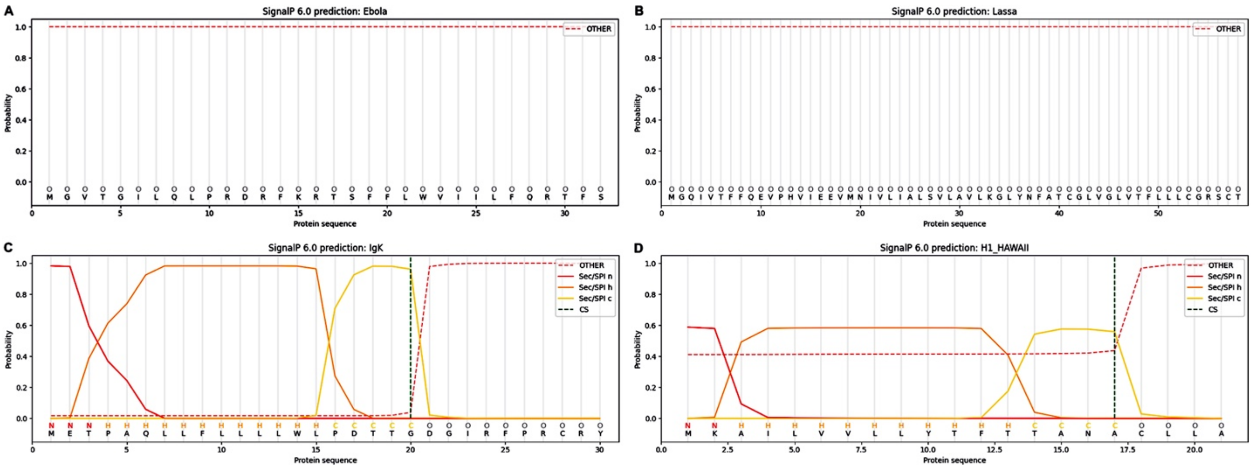

**Supplementary Figure 2. Validation of the structure-based prediction of SPs using the bacterial *Yersinia pestis* SP. (A)** Prediction of the bacterial *Y. pestis* SP using signalP6 bacterial mode. **(B)** Structural model of the bacterial *Y. pestis* SP in complex with the ffh protein, the equivalence of the human SRP54 in bacteria. **(C)** Structural model of the bacterial *Y. pestis* SP in complex with the human SRP54. **(D)** Predicted SP regions of the bacterial *Y. pestis* SP through structural modeling.

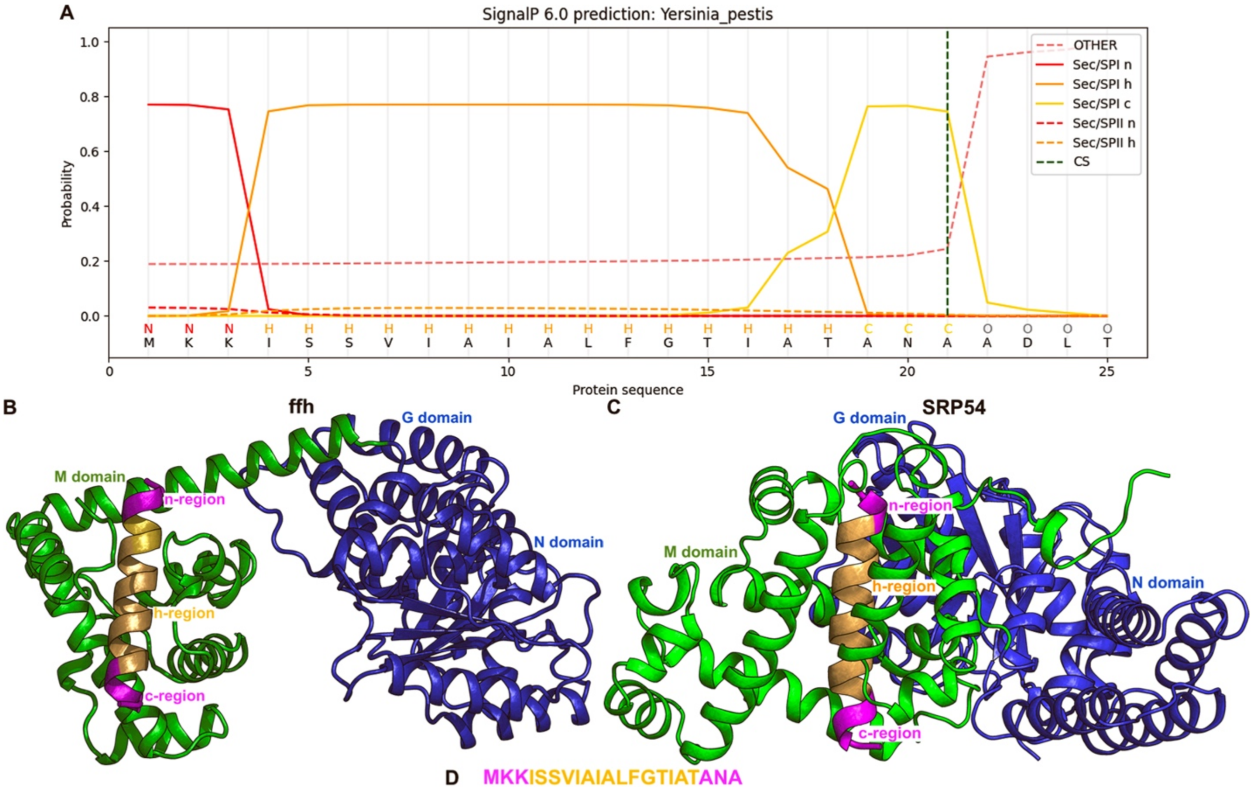

**Supplementary Figure 3. Structural models of the bacterial *Y. pestis* SP in complex with the bacterial signal peptidase 1 and human SPC-C.** The models illustrated the cleavage site at the c-region of the *Y. pestis* SP by the signal peptidases.

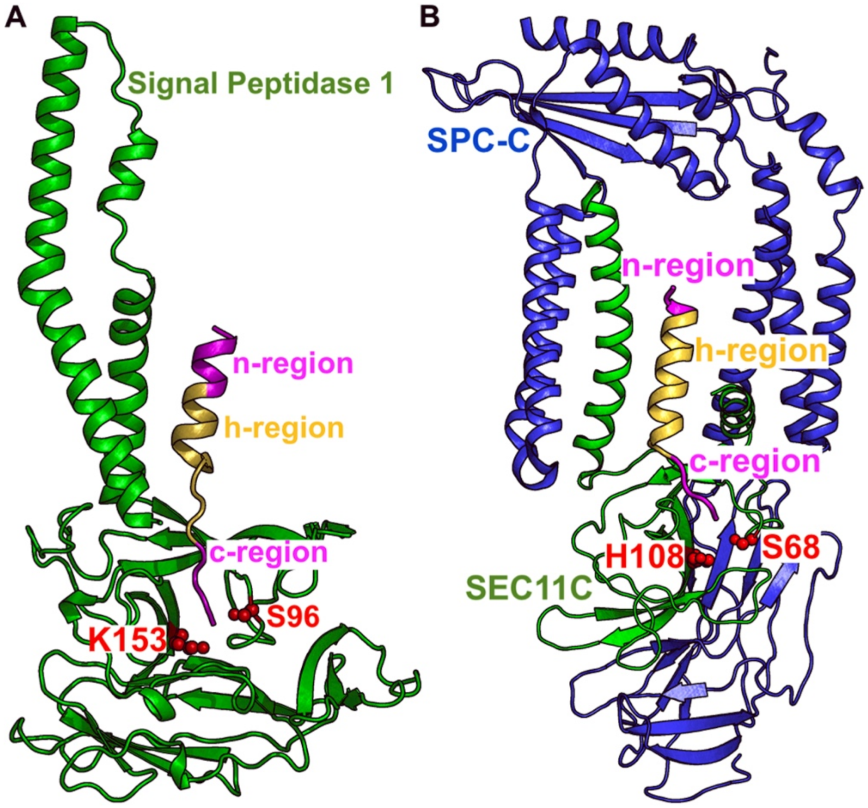

**Supplementary Figure 4. Structural models of the SPs of Ebola and Lassa glycoproteins as well as Igκ and H1 HAWAII in complex with the human SPC-C.** The models illustrated the cleavage sites at the c-regions of the SPs and supported the findings in **Figure 2**.

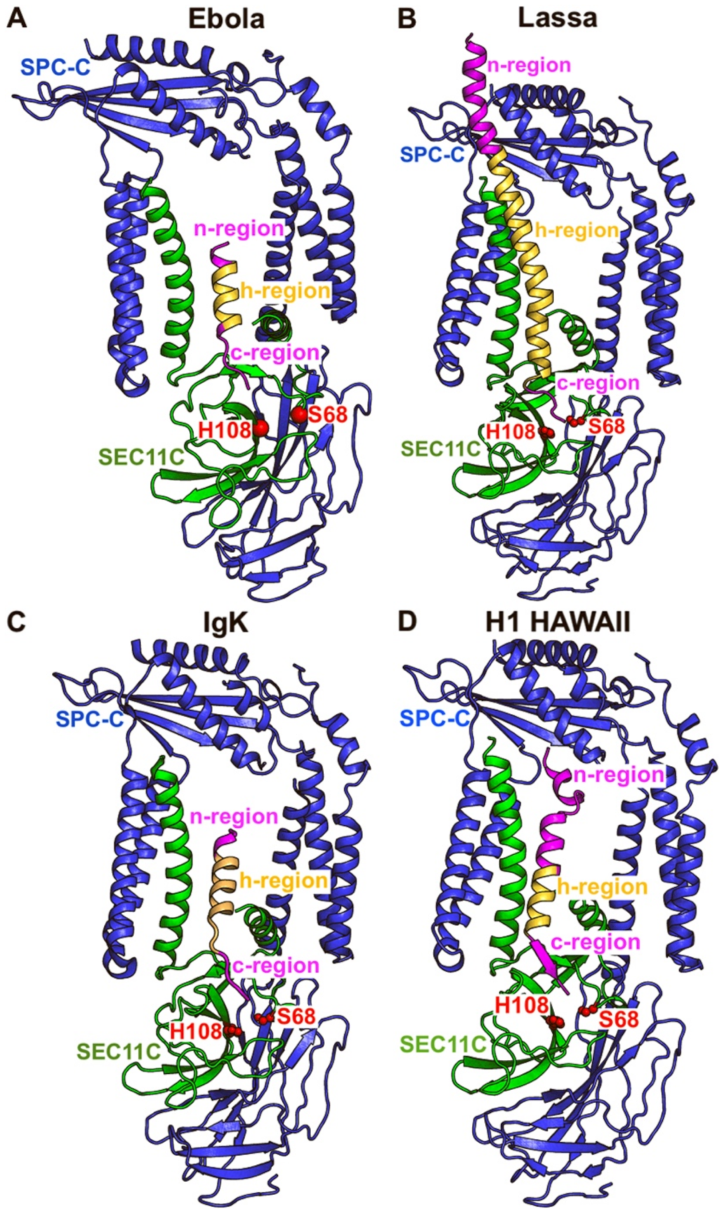

**Supplementary Figure 5. Plots of the hydrophobicity of the Ebola, Lassa, Igκ, and H1 HAWAII, and bacterial *Y. pestis* SPs calculated based on the Kyte-Doolittle[1] hydrophobicity scale.** The n-, h-, and c-regions of SPs were labeled in the plot, with the total and average hydrophobicity of the n, h, and c-regions included in the center of the corresponding panels.

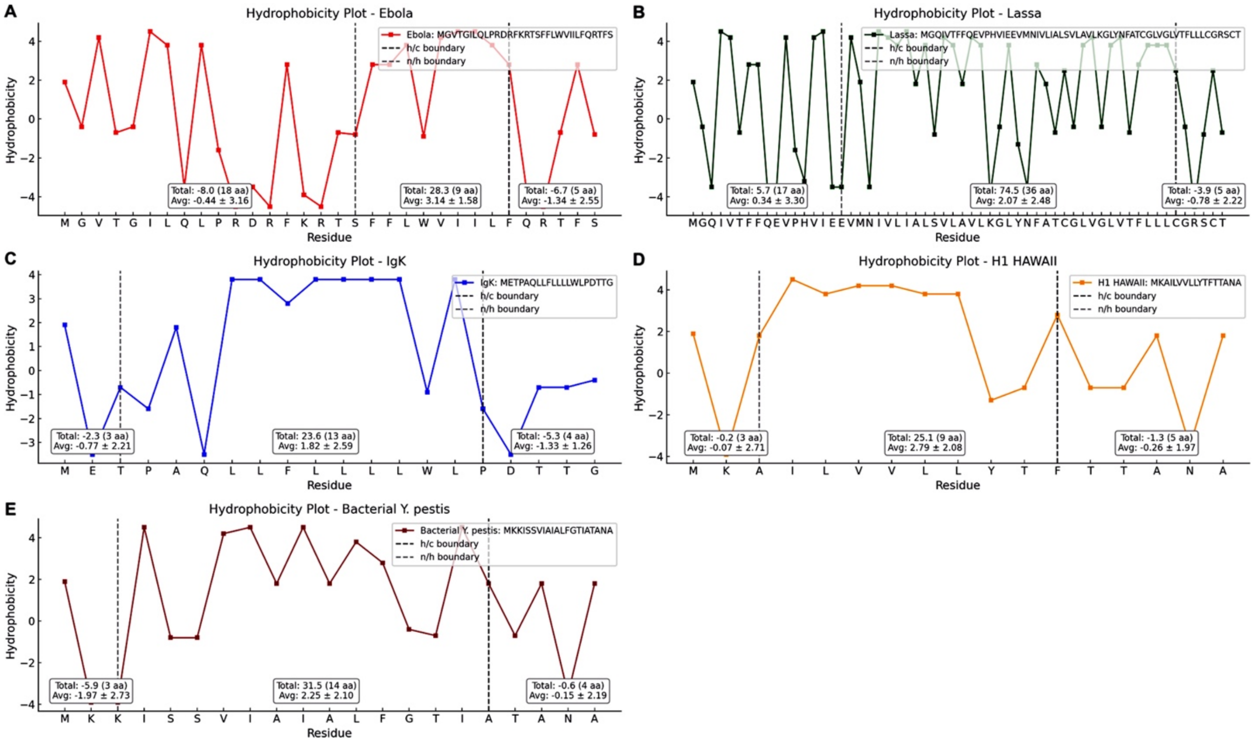

**Supplementary Figure 6. Evidence for the proposed computational workflow for structure-based optimization of non-functional SPs using the human serum albumin (HSA) and H1 HAWAII SPs**. Structural model of the H1 HAWAII SP in complex with the ER lipids **(A)** were included here, while the structural models of the H1 HAWAII SP in complex with SRP54 and SPC-C could be found in **Figure 2D** and **Supplementary Figure 4D**. Structural models of the HSA SP in complex with the ER lipids **(B)**, SRP54 **(C)**, and SPC-C **(D)** were built and shown in this figure. The predicted binding energy of the complexes could be found in **Supplementary Table 2**.

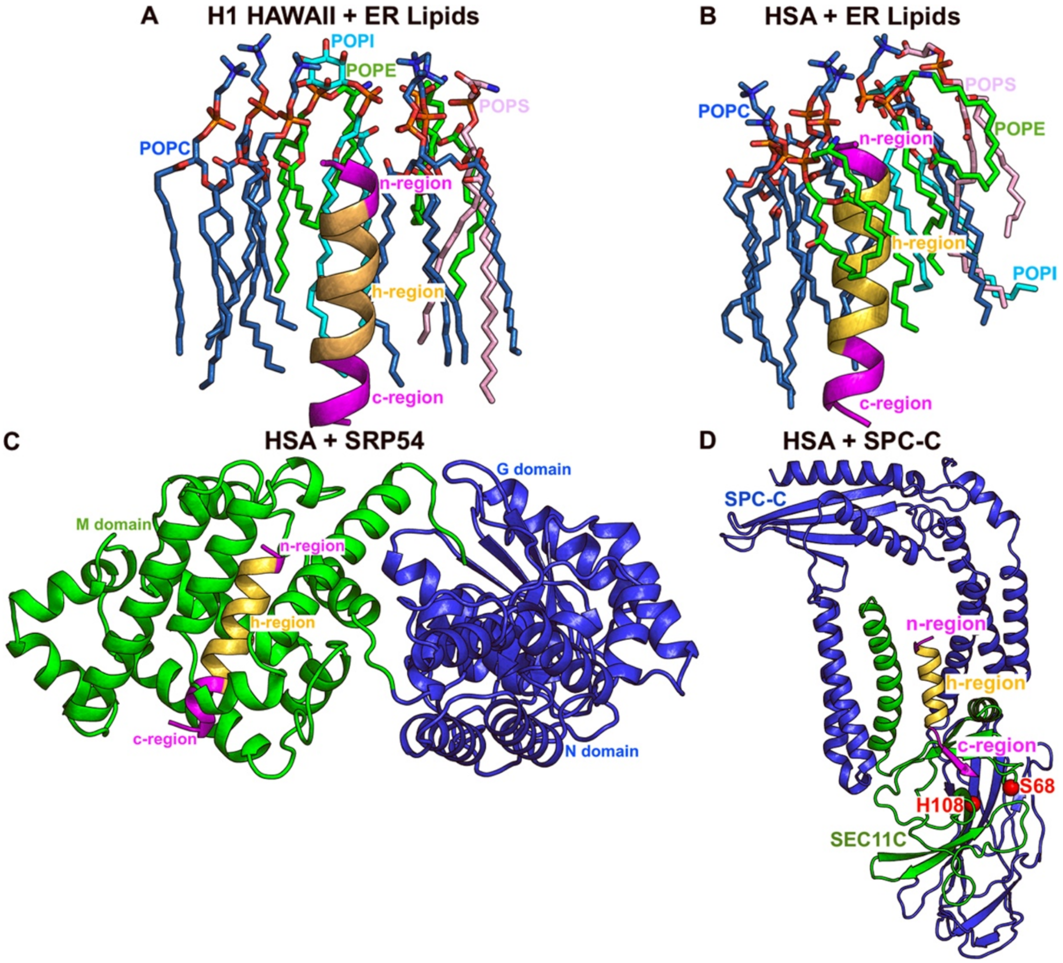

**Supplementary Figure 7. Plot of hydrophobicity of the HSA SP calculated based on the Kyte-Doolittle[1] hydrophobicity scale.** The n-, h-, and c-regions of SP were labeled in the plot, with the total and average hydrophobicity of the n-, h-, and c-regions included in the center of the corresponding panels.

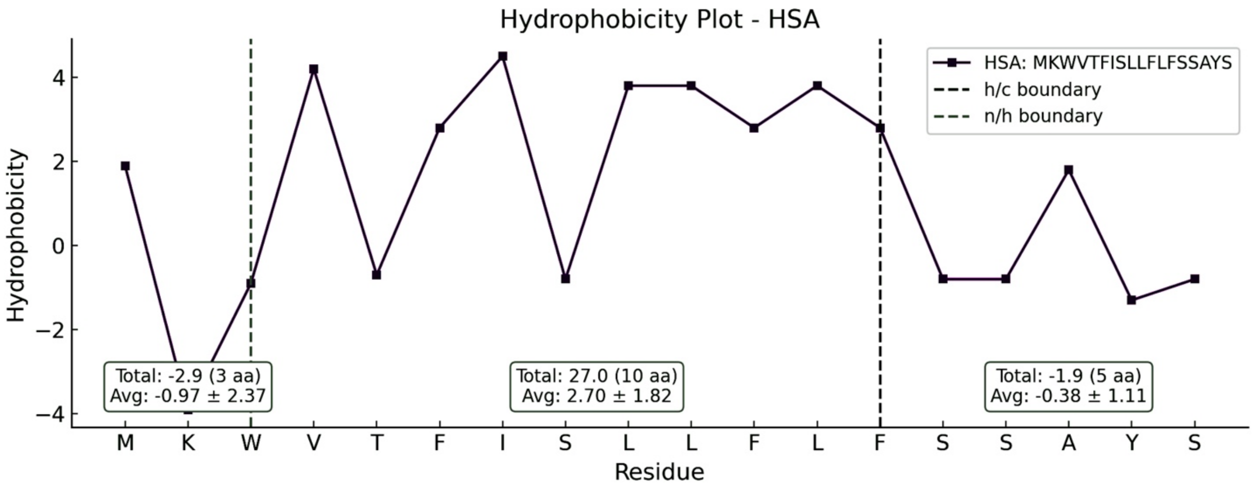

**Supplementary Figure 8. Structural models of the bacterial *Y. pestis* SP in complex with the ER lipids and of the optimized *Y. pestis* SP (using the most confident redesigned sequence by LigandMPNN[2,3]) for protein expression in human cells in complex with the ER lipids, human SRP54, and SPC-C.** The models of bacterial *Y. pestis* SP in complex with human SRP54 and SPC-C can be found in **Figures S2C** and **Supplementary Figure 3B**, respectively. The predicted binding energy of the complexes could be found in **Supplementary Table 3**.

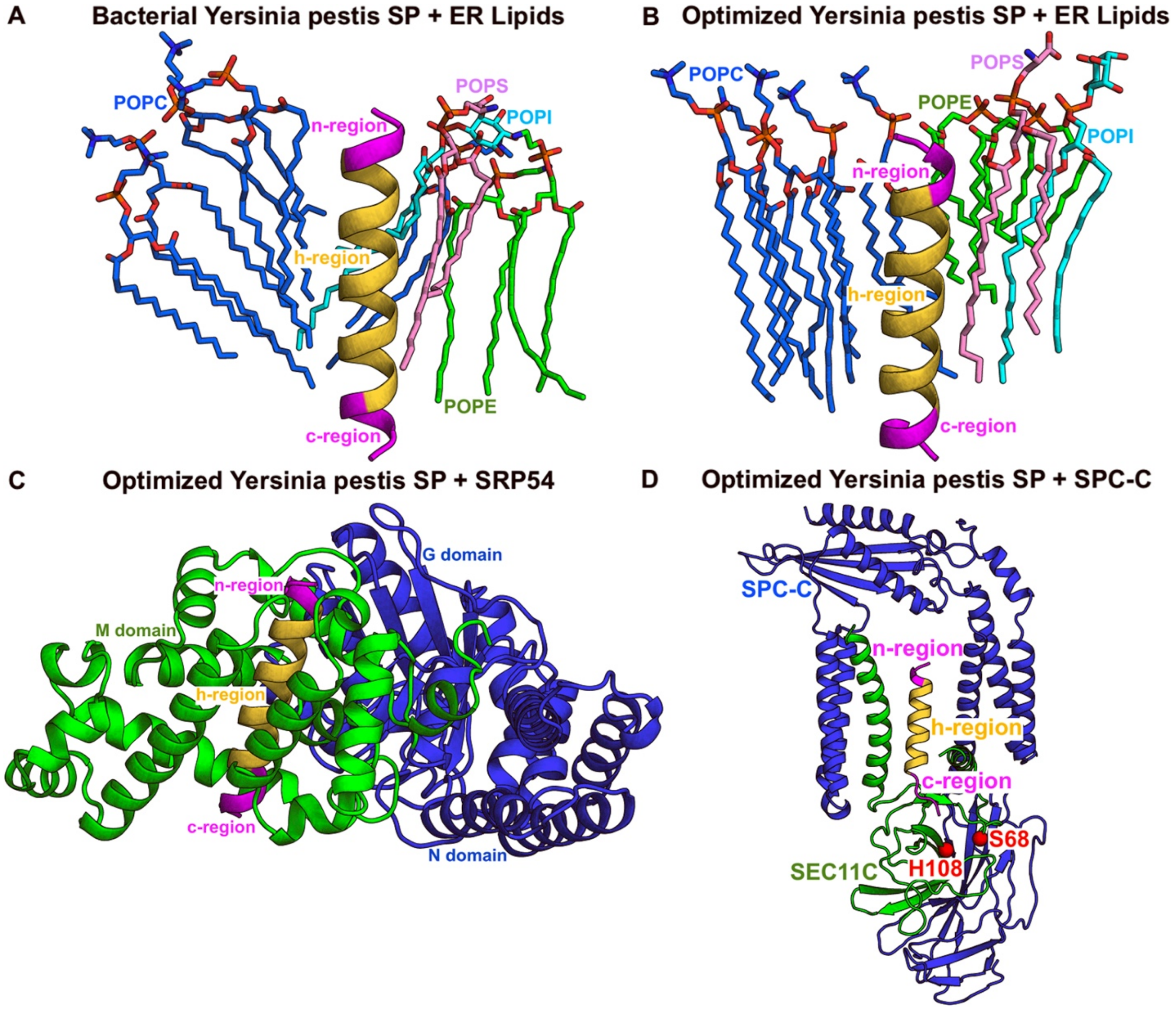

**Supplementary Figure 9. Plot of hydrophobicity of the optimized *Y. pestis* SP (using the most confident redesigned sequence by LigandMPNN[2,3]) calculated based on the Kyte-Doolittle[1] hydrophobicity scale.** The n-, h-, and c-regions of SP were labeled in the plots, with the total and average hydrophobicity of the h-region included in the center panels.

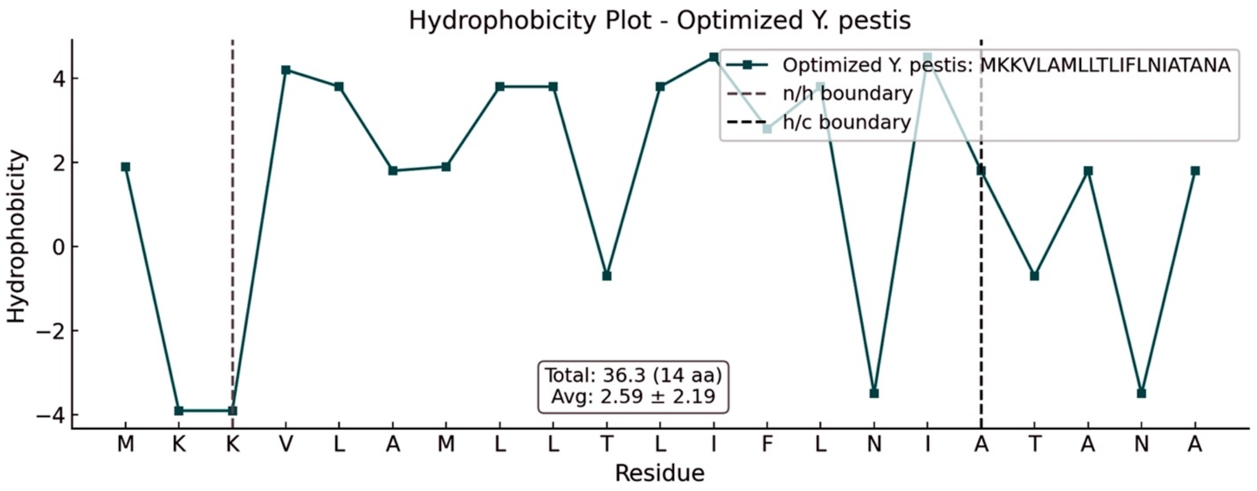

**Supplementary Figure 10. Structural models of the Lassa SPs in complex with its glycoprotein complex (GPC), ER lipids and SPC-C.** The structural model of Lassa SP in complex with SRP54 can be found in **Figure 2B**. The predicted binding energy of the complexes could be found in **Supplementary Table 5.**

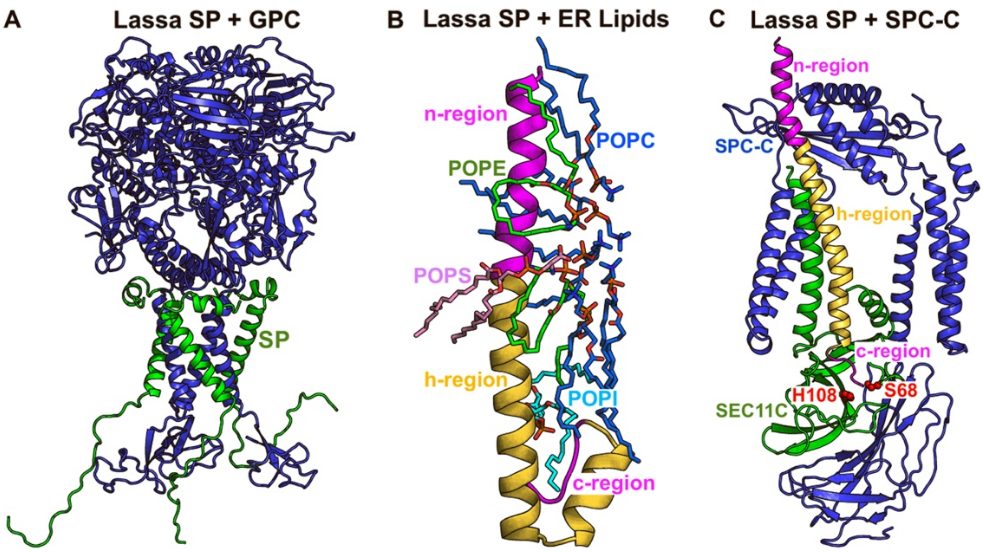

**Supplementary Figure 11. Structural models of the optimized Lassa SPs (using the most confident redesigned sequence by LigandMPNN[2,3]) in complex with its glycoprotein complex (GPC), ER lipids, SRP54, and SPC-C.** The predicted binding energy of the complexes could be found in **Supplementary Table 5.**

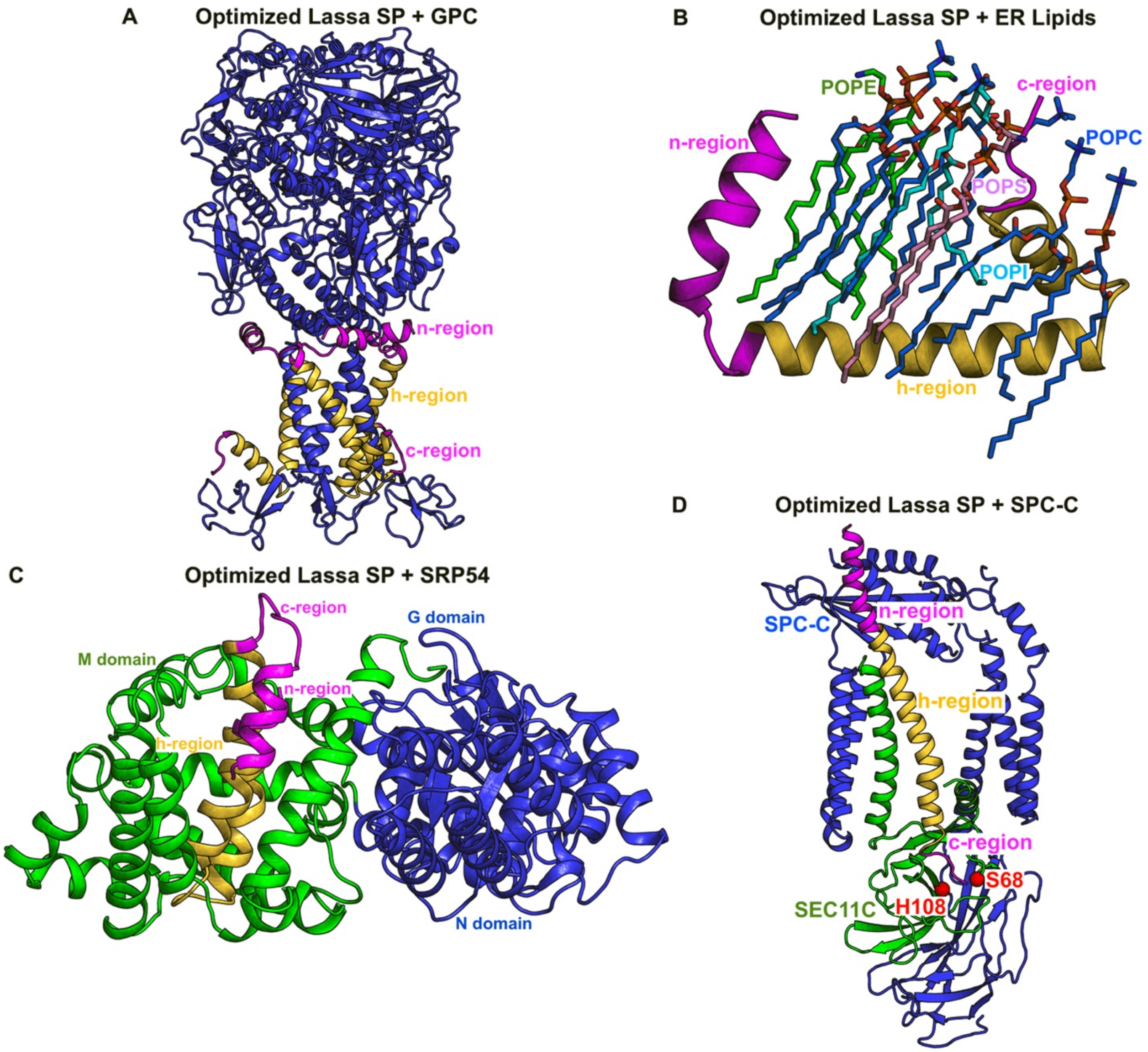

**Supplementary Figure 12. Plot of hydrophobicity of the optimized Lassa SP (using the most confident redesigned sequence by LigandMPNN[2,3]) calculated based on the Kyte-Doolittle[1] hydrophobicity scale.** The n-, h-, and c-regions of SP were labeled in the plot, with the total and average hydrophobicity of the h-region included in the center panel.

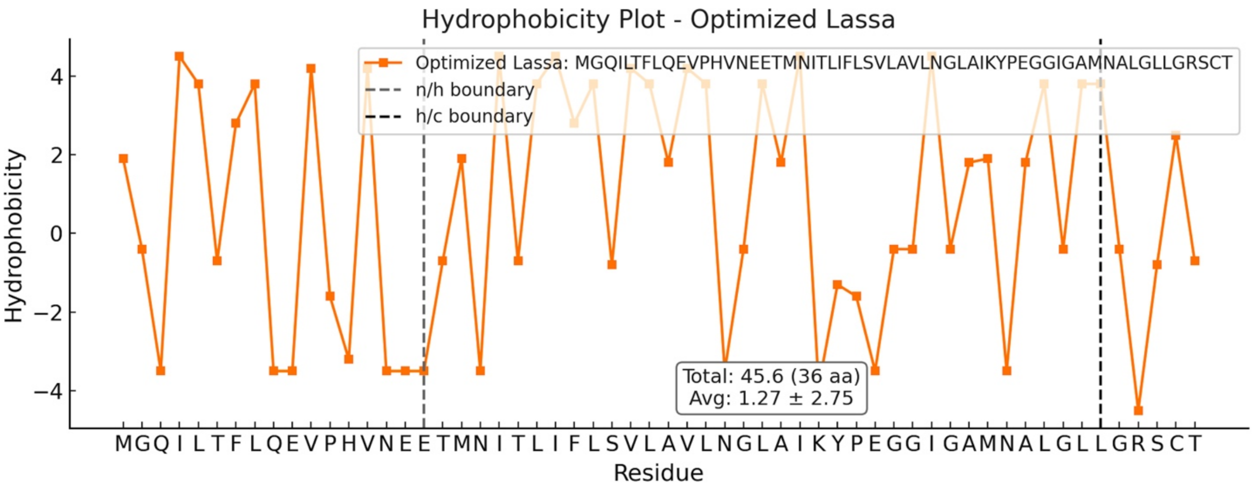

**Supplementary Figure 13. Structural models of the VEEV E3 protein (SLVTTMCLLANVTFPCAQPPICYDRKPAETLAMLSVNVDNPGYDELLEAAVKCPGRKRR), which serves as the SP of VEEV, in complex with its GPC, ER lipids, SRP54, and SPC-C.** VEEV E3 was not found in the SEC11C catalytic pocket of SPC-C, given the absence of cleavage motif. The predicted binding energy of the complexes could be found in **Supplementary Table 7.**

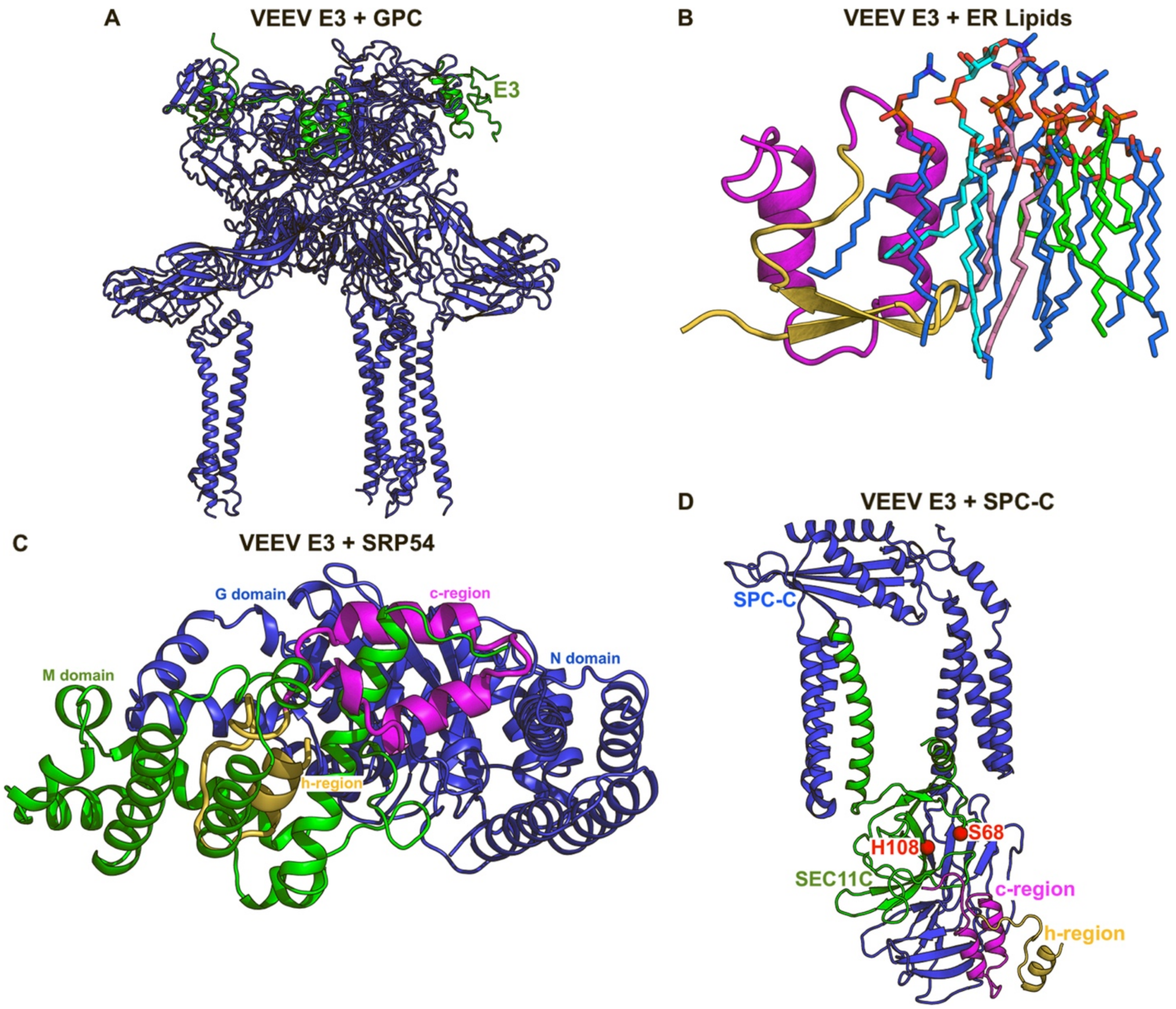

**Supplementary Figure 14. Structural models of the optimized VEEV E3 protein (using the most confident redesigned sequence by LigandMPNN[2,3]) in complex with its GPC, ER lipids, SRP54, and SPC-C.** The optimized VEEV E3 was not found in the SEC11C catalytic pocket of SPC-C. The predicted binding energy of the complexes could be found in **Supplementary Table 7.**

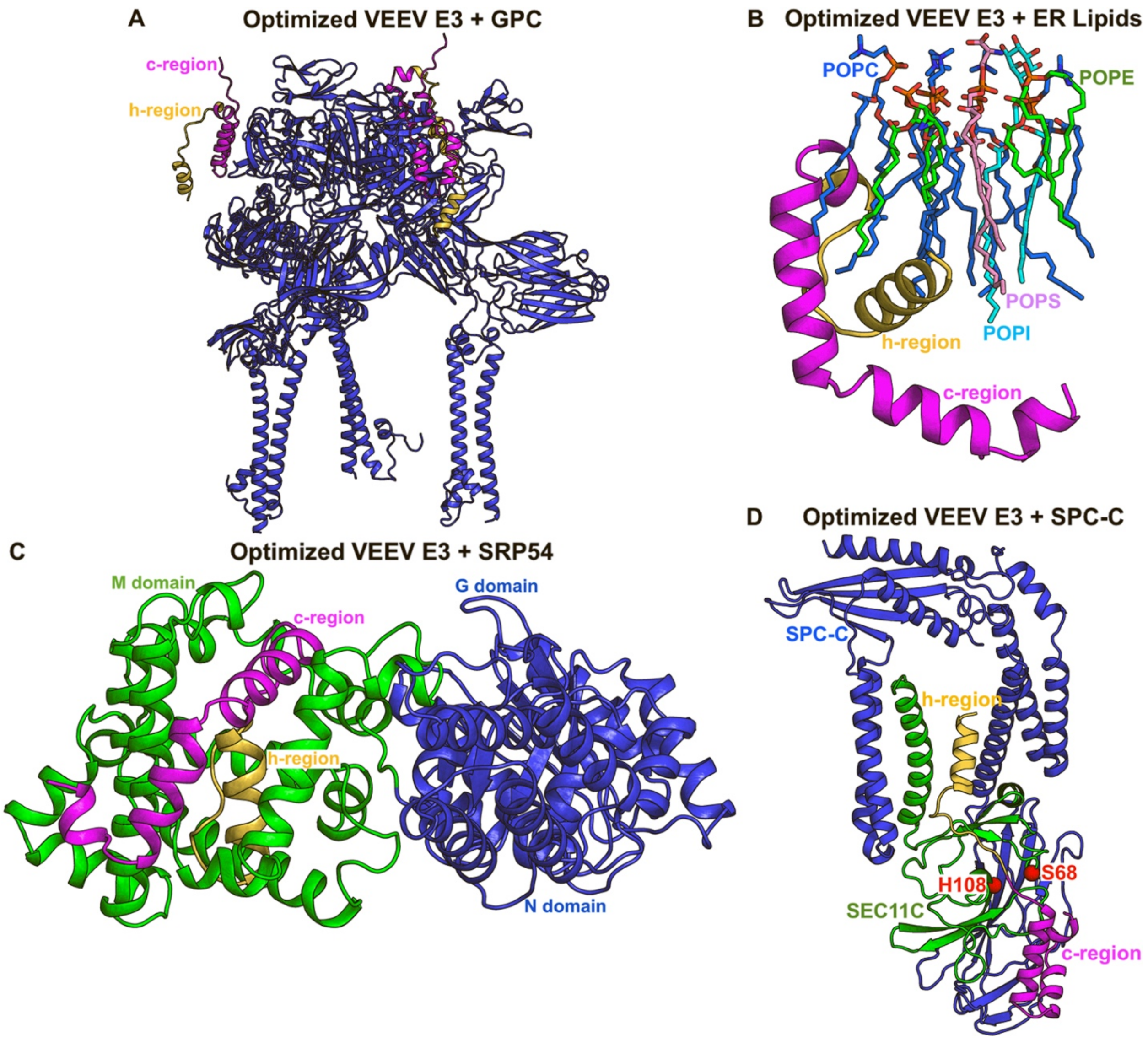

**Supplementary Figure 15. Plots of hydrophobicity of the native and optimized VEEV SP (using the most confident redesigned sequence by LigandMPNN[2,3]) calculated based on the Kyte-Doolittle[1] hydrophobicity scale.** The n-, h-, and c-regions of SPs were labeled in the plots, with the total and average hydrophobicity of the h-regions included in the center panels.

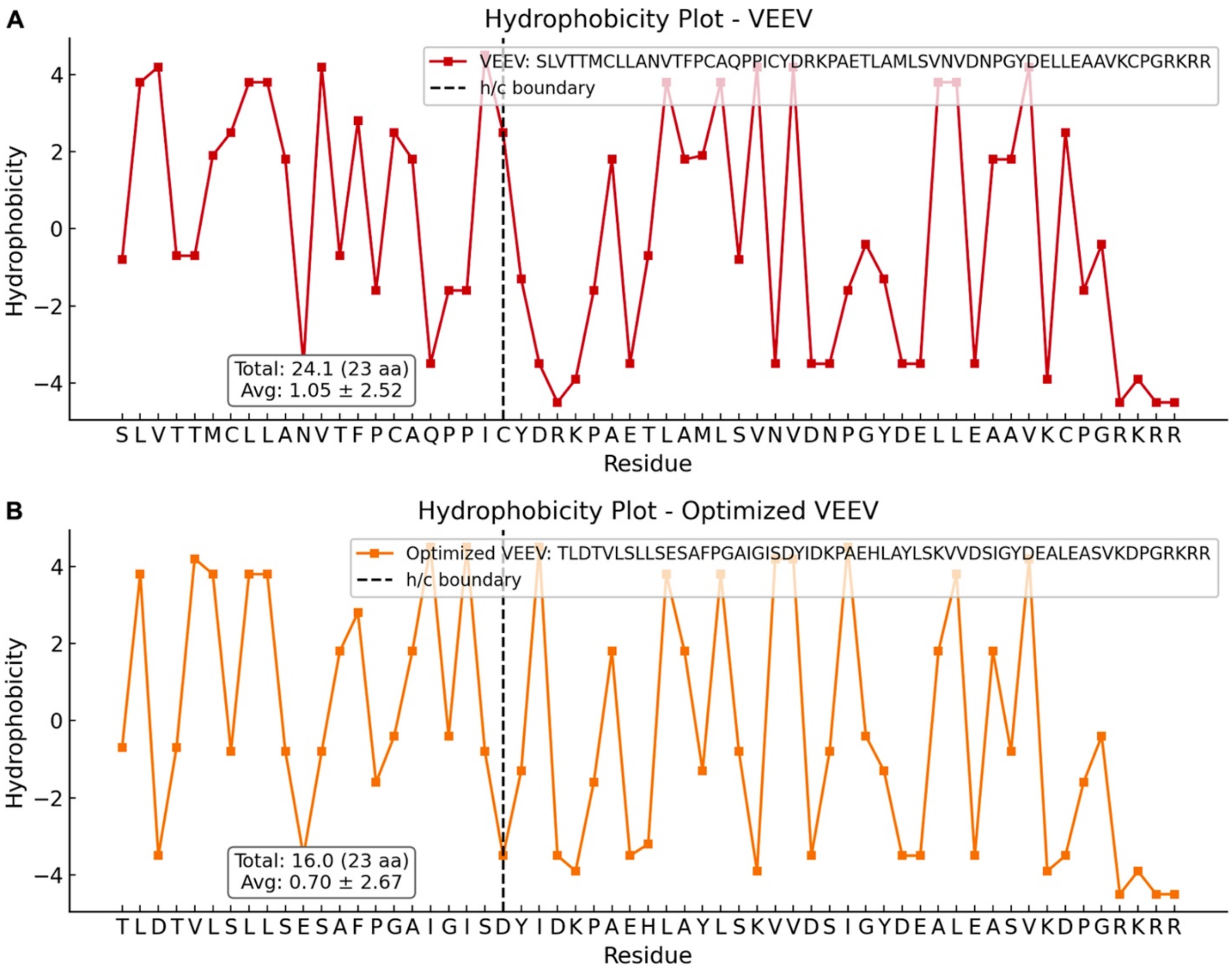

**Supplementary Figure 16. Cellular protein expression following F1 mRNA transfection. A, B.** Western blot analysis of *Y. pestis* F1 protein with different SPs or mock transfected samples as noted above the gel images; Lane designations: L, molecular weight ladder; 1, cell supernatant fraction; 2, cytosolic fraction; 3, cell membrane fraction; +, positive control, **C**. Protein expression levels for each fraction measured from the Western blot images, and **D**. Total protein expression levels for the four F1 mRNA constructs.

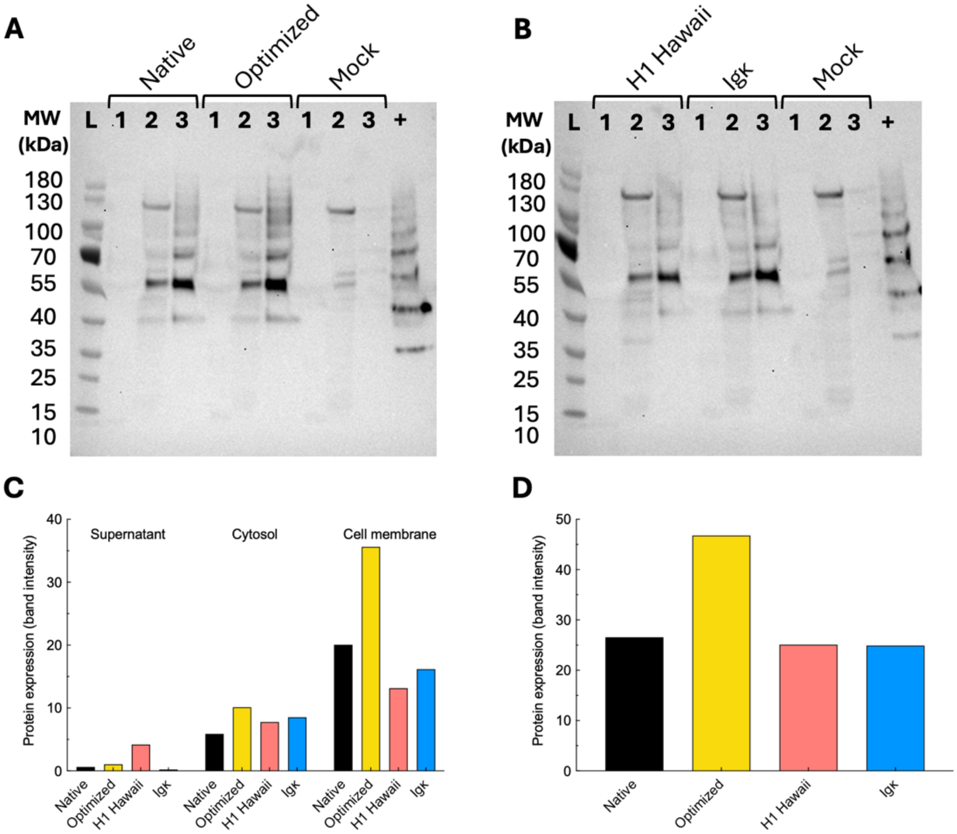

**Supplementary Figure 17. Comparison of the proportions of helix-favoring and helix-breaking residues in the h-regions of H1 HAWAII versus HSA, bacterial versus optimized *Y. pestis* SP, native versus redesigned Lassa SP, and native versus optimized VEEV E3 protein**. The secondary structure propensities of the amino acids were based on the Chou-Fasman amino acid propensities[4]. The list of helix-favoring residues included A, E, L, M, Q, K, R, and H, while the helix-breaking residues were P, G, Y, N, S, and D[4].

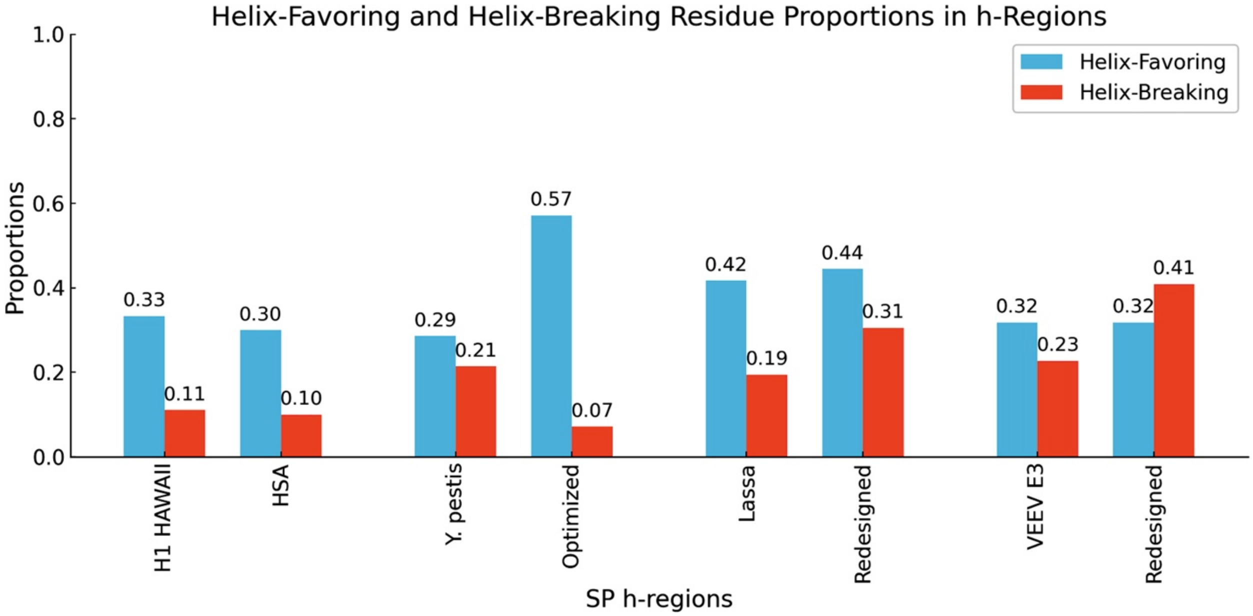

**Supplementary Table 1. Predicted binding energy of the H1 HAWAII and HSA SPs to the ER lipids, SRP54, and SPC-C**. The binding energy between the SPs and ER lipids were measured using PRODIGY-LIG[5,6], while between the SPs and SRP54 or SPC-C were determined using PRODIGY[7,8], whose outputs included the number of interactions between the two proteins.

| SP | Binding Partner | ΔG(kcal/mol) | Number of Interactions | ΔG(kcal/mol/ interaction) |
|---|---|---|---|---|
| H1 HAWAII | ER Lipids | -53.0 | - | - |
| | SRP54 | -10.9 | 111 | -0.098 |
| | SPC-C | -9.2 | 69 | -0.13 |
| HSA | ER Lipids | -56.5 | - | - |
| | SRP54 | -11.1 | 111 | -0.1 |
| | SPC-C | -10.0 | 71 | -0.14 |

**Supplementary Table 2. Top 10 sequences of the optimized *Y. pestis* SP by applying LigandMPNN[2,3] on the structural model of bacterial *Y. pestis* SP in complex with human SRP54 to redesign the SP in Supplementary Figure 2C.**

| **Redesigned *Y. pestis* SP** | **LigandMPNN Confidence Score** |
|---|---|
| MKKVLAMLLTTIFLNIATANA | 0.4604 |
| MKKVLAMLLTTIFLNIATANA | 0.4732 |
| MKKVLAMLLLTIFCNIATANA | 0.4267 |
| MKKVLAMLLTLIFLNIATANA | 0.4780 |
| MKKVLAMLLTTIFLNIATANA | 0.4608 |
| MKKVLAMLLTTIFLNIATANA | 0.4727 |
| MKKVLAMLLTTILLNIATANA | 0.4544 |
| MKKVLAMLLTTILLNIATANA | 0.4596 |
| MKKVLAMLLTTIFLNIATANA | 0.4660 |
| MKKVLAMLLLTIFCNIATANA | 0.4391 |

**Supplementary Table 3. Predicted binding energy of the bacterial and optimized *Y. pestis* SPs to the ER lipids, SRP54, and SPC-C.** The binding energy between the SPs and ER lipids were measured using PRODIGY-LIG[5,6], while between the SPs and SRP54 or SPC-C were determined using PRODIGY[7,8], whose outputs included the number of interactions between the two proteins.

| SP | Binding Partner | ΔG(kcal/mol) | Number of Interactions | ΔG(kcal/mol/ interaction) |
|---|---|---|---|---|
| Bacterial *Y. pestis* | ER Lipids | -54.8 | - | - |
| | SRP54 | -10.0 | 113 | -0.088 |
| | SPC-C | -9.5 | 80 | -0.12 |
| Optimized *Y. pestis* | ER Lipids | -56.1 | - | - |
| | SRP54 | -12.4 | 121 | -0.1 |
| | SPC-C | -8.7 | 65 | -0.14 |

**Supplementary Table 4. Top 10 sequences of the optimized Lassa viral SP by applying LigandMPNN[2,3] on the structural model of Lassa viral SP in complex with human SRP54 to redesign the SP with the residues involved in the interactions with its GPC kept fixed.**

| Redesigned Lassa SP | LigandMPNN Confidence Score |
|---|---|
| MGQILTFLQEVPHVNEETMNITLIFLSVLAVLNGLAIKYPEGGIGAMNALGLLGRSCT | 0.3375 |
| MGQILTFLQEVPHVNEEVMNITLILLSVLAVGYGLAVKYPEGGVGFIAALGLLGRSCT | 0.3288 |
| MGQILTFLQEVPHVNEEVMNITLILLSVLAVAYGLAVKYPEGGIGMMNAFGLLGRSCT | 0.3086 |
| MGQILTFLQEVPHVNEEVMNITLIFLSVLAVLNGLGIKSPEGAIGAINALGLLGRSCT | 0.3165 |
| MGQILTFLQEVPHVNEEVMNITLILLSVLAVGYGLGVKSPEGLIGMINALGLLGRSCT | 0.3199 |
| MGQILTFLQEVPHVNEETMNIVLIFLSVLAVGNGLAIKYPDGGIGFNAALGLLGRSCT | 0.3099 |
| MGQILTFLQEVPHVNEETMNITLILLSVLAVAYGLGVKSPEGLIGMINAFGLMGRSCT | 0.3269 |
| MGQILTFLQEVPHVNEEVMNITLIFLSVLAVLYGLAVKYPEGGIGAINALGLLGRSCT | 0.3360 |
| MGQILTFLQEVPHVVEETMNITLILLSVLAVLNGLAIKYPEGGIGAINALGLLGRSCT | 0.3350 |
| MGQILTFLQEVPHVNEEVMNITLILLSVLAVLRGLAIKYPEGGIGAINALGQLGRSCT | 0.3201 |

**Supplementary Table 5. Predicted binding energy of the native and optimized Lassa SPs to its glycoprotein complex (GPC), ER lipids, SRP54, and SPC-C.** The binding energy between the SPs and ER lipids were measured using PRODIGY-LIG[5,6], while between the SPs and SRP54 or SPC-C were determined using PRODIGY[7,8], whose outputs included the number of interactions between the two proteins.

| SP | Binding Partner | ΔG(kcal/mol) | Number of Interactions | ΔG(kcal/mol/ interaction) |
|---|---|---|---|---|
| Lassa | GPC | -8.2 | 97 | -0.085 |
| | ER Lipids | -62.8 | - | - |
| | SRP54 | -14.6 | 183 | -0.080 |
| | SPC-C | -14.1 | 146 | -0.097 |
| Optimized Lassa | GPC | -13.5 | 133 | -0.10 |
| | ER Lipids | -61.9 | - | - |
| | SRP54 | -18.9 | 221 | -0.086 |
| | SPC-C | -13.2 | 137 | -0.096 |

**Supplementary Table 6. Top 10 sequences of the optimized VEEV E3 protein by applying LigandMPNN[2,3] on the structural model of VEEV E3 in complex with human SRP54 to redesign the SP h-region with the residues involved in the interactions with its GPC kept fixed.**

| Redesigned VEEV E3 | LigandMPNN Confidence Score |
|---|---|
| TLGTALSLLSESAFPGAISIADYIDKPAETLAYLSKVVDSLGYDELLEASVKDPGRKRR | 0.3867 |
| TLDTVLSLMSEGAFPGAIGITDYIENPAELLAYLSKKVDDIGYDELLEASVKDPGRKRR | 0.3802 |
| TLDTVLSLLSESAFPGAIGISDYIDKPAEHLAYLSKVVDSIGYDEALEASVKDPGRKRR | 0.3972 |
| TLGTVLSLLSDSRFPGAISIFDYIDNPAEHLAYLSKVVDSIGYDEALEASVKFEGRKRR | 0.3913 |
| TLGTALSLMSESRFPGAISIFDYIENPAETLAYLSKVVDSIGYDELLEACVKDPGRKRR | 0.3879 |
| TLGTVLSLLSESKFPGAISIFDYEDNPAEHLAYLSKVVDSIGYDEALEASVKDPGRKRR | 0.3783 |
| TLDTALSLLSEGKFPGALGITHYEENPAEHLAYLSKVVDSLGYDEALEASVKDEGRKRR | 0.3649 |
| TLGTVLSLMSETRFPGAISISDYIDNPAETLAYLSKVVDSIGYDEFLEASVKDPGRKRR | 0.3759 |
| TLGTVLSLLSESAFPGAISIFDYIDNPAETLAYLSKVVDSVGYDELLEACVKDPGRKRR | 0.3970 |
| TLGTVLSLLSETAFPGAISIFDYIDNPAETLAYLSKNVDSIGYDELLEASVKFEGRKRR | 0.3870 |

**Supplementary Table 7. Predicted binding energy of the native and optimized VEEV E3 protein to its GPC, ER lipids, SRP54, and SPC-C.** The binding energy between the SPs and ER lipids were measured using PRODIGY-LIG[5,6], while between the SPs and SRP54 or SPC-C were determined using PRODIGY[7,8], whose outputs included the number of interactions between the two proteins.

| SP | Binding Partner | ΔG(kcal/mol) | Number of Interactions | ΔG(kcal/mol/ interaction) |
|---|---|---|---|---|
| VEEV E3 | GPC | -6.2 | 57 | -0.11 |
| | ER Lipids | -51.4 | - | - |
| | SRP54 | -16.5 | 190 | -0.085 |
| | SPC-C | -6.2 | 30 | -0.21 |
| Optimized VEEV E3 | GPC | -5.4 | 53 | -0.10 |
| | ER Lipids | -48.8 | - | - |
| | SRP54 | -18.2 | 209 | -0.087 |
| | SPC-C | -12.2 | 141 | -0.087 |

**Supplementary Data 1. Sequences of the pathogenic proteins used in our experimental validations.**

***Y. pestis*** **F1 - Native (SP sequence is underlined)**

MKKISSVIAIALFGTIATANAADLTASTTATATLVEPARITLTYKEGAPITIMDNGNIDTELLVGTLTLGGYKTGTTSTSVNFTDAAGDPMYLTFTSQDGNNHQFTTKVIGKDSRDFDISPKVNGENLVGDDVVLATGSQDFFVRSIGSKGGKLAAGKYTDAVTVTVSNQ

***Y. pestis*** **F1 – with H1 Hawaii SP (SP sequence is underlined)**

MKAILVVLLYTFTTANAKKISSVIAIALFGTIATANAADLTASTTATATLVEPARITLTYKEGAPITIMDNGNIDTELLVGTLTLGGYKTGTTSTSVNFTDAAGDPMYLTFTSQDGNNHQFTTKVIGKDSRDFDISPKVNGENLVGDDVVLATGSQDFFVRSIGSKGGKLAAGKYTDAVTVTVSNQ

***Y. pestis*** **F1 – with Igκ SP (SP sequence is underlined)**

METPAQLLFLLLLWLPDTTGKKISSVIAIALFGTIATANAADLTASTTATATLVEPARITLTYKEGAPITIMDNGNIDTELLVGTLTLGGYKTGTTSTSVNFTDAAGDPMYLTFTSQDGNNHQFTTKVIGKDSRDFDISPKVNGENLVGDDVVLATGSQDFFVRSIGSKGGKLAAGKYTDAVTVTVSNQ

***Y. pestis*** **F1 – with optimized SP (SP sequence is underlined)**

MKKVLAMLLLLIFLVIATANAADLTASTTATATLVEPARITLTYKEGAPITIMDNGNIDTELLVGTLTLGGYKTGTTSTSVNFTDAAGDPMYLTFTSQDGNNHQFTTKVIGKDSRDFDISPKVNGENLVGDDVVLATGSQDFFVRSIGSKGGKLAAGKYTDAVTVTVSNQ

**Lassa GPC – with optimized SP (SP sequence is underlined)**

MGQILTFLQEVPHVNEETMNITLIFLSVLAVLNGLAIKYPEGGIGAMNALGLLGRSCTTSLYKGVYELQTLELNMETLNMTMPLSCTKNNSHHYIMVGNETGLELTLTNTSIINHKFCNLSDAHKKNLYDHALMSIISTFHLSIPNFNQYEAMSCDFNGGKISVQYNLSHSYAGDAANHCGTVANGVLQTFMRMAWGGSYIALDSGRGNWDCIMTSYQYLIIQNTTWEDHCQFSRPSPIGYLGLLSQRTRDIYISRRLLGTFTWTLSDSEGKDTPGGYCLTRWMLIEAELKCFGNTAVAKCNEKHDEEFCDMLRLFDFNKQAIQRLKAEAQMSIQLINKAVNALINDQLIMKNHLRDIMGIPYCNYSKYWYLNHTTTGRTSLPKCWLVSNGSYLNETHFSDDIEQQADNMITE

MLQKEYMERQGKTPLGLVDLFVFSTSFYLISIFLHLVKIPTHRHIVGKSCPKPHRLNHMGICSCGLYKQPGVPVKWKR

**VEEV E3/E2/6k/E1 – with optimized SP (SP sequence is underlined)**

MTLDTVLSLLSESAFPGAIGISDYIDKPAEHLAYLSKVVDSIGYDEALEASVKDPGRKRRSTEELFKEYKLTRPYMARCIRCAVGSCHSPIAIEAVKSDGHDGYVRLQTSSQYGLDSSGNLKGRTMRYDMHGTIKEIPLHQVSLHTSRPCHIVDGHGYFLLARCPAGDSITMEFKKDSVTHSCSVPYEVKFNPVGRELYTHPPEHGVEQACQVYAHDAQNRGAYVEMHLPGSEVDSSLVSLSGSSVTVTPPVGTSALVECECGGTKISETINKTKQFSQCTKKEQCRAYRLQNDKWVYNSDKLPKAAGATLKGKLHVPFLLADGKCTVPLAPEPMITFGFRSVSLKLHPKNPTYLTTRQLADEPHYTHELISEPAVRNFTVTEKGWEFVWGNHPPKRFWAQETAPGNPHGLPHEVITHYYHRYPMSTILGLSICAAIATVSVAASTWLFCRSRVACLTPYRLTPNARIPFCLAVLCCARTARAETTWESLDHLWNNNQQMFWIQLLIPLAALIVVTRLLRCVCCVVPFLVMAGAAGAGAYEHATTMPSQAGISYNTIVNRAGYAPLPISITPTKIKLIPTVNLEYVTCHYKTGMDSPAIKCCGSQECTPTYRPDEQCKVFTGVYPFMWGGAYCFCDTENTQVSKAYVMKSDDCLADHAEAYKAHTASVQAFLNITVGEHSIVTTVYVNGETPVNFNGVKLTAGPLSTAWTPFDRKIVQYAGEIYNYDFPEYGAGQPGAFGDIQSRTVSSSDLYANTNLVLQRPKAGAIHVPYTQAPSGFEQWKKDKAPSLKFTAPFGCEIYTNPIRAENCAVGSIPLAFDIPDALFTRVSETPTLSAAECTLNECVYSSDFGGIATVKYSASKSGKCAVHVPSGTATLKEAAVELTEQGSATIHFSTANIHPEFRLQICTSYVTCKGDCHPPKDHIVTHPQYHAQTFTAAVSKTAWTWLTSLLGGSAVIIIIGLVLATIVAMYVLTNQKHN
